# Increased public health threat of avian-origin H3N2 influenza virus during evolution in dogs

**DOI:** 10.1101/2022.10.10.511550

**Authors:** Mingyue Chen, Yanli Lyu, Fan Wu, Ying Zhang, Hongkui Li, Rui Wang, Yang Liu, Xinyu Yang, Liwei Zhou, Ming Zhang, Qi Tong, Honglei Sun, Juan Pu, Jinhua Liu, Yipeng Sun

## Abstract

Influenza A viruses in animal reservoirs repeatedly cross species barrier to infect humans. Once an animal-borne virus with novel antigenicity acquired the efficient human to human transmissibility, it will become epidemic in the population. Dogs are the closest animal companions to humans and canine respiratory tract expresses both SAα2,3-(avian type) and α2,6-Gal (human type) receptors. However, the role of dogs in the ecology of influenza viruses is unclear. H3N2 avian influenza viruses transmitted to dogs around 2006 and have formed stable lineages. The long-term epidemic of avian-origin H3N2 virus in canine offers the best models to investigate the effect of dogs on the evolution of influenza viruses. Here, we carried out a systematic and comparative identification of the biological characteristics of H3N2 canine influenza viruses (CIVs) isolated in the worldwide over 10 years. We found that during the adaptation of H3N2 CIVs to dogs, H3N2 CIVs became to recognize the human-like SAα2,6-Gal receptor, gradually increased HA acid stability and replication ability in human airway epithelial cells, and acquired a 100% transmission rate via respiratory droplet in ferret model, which were essential hallmarks of being adapted to humans. We also identified that the frequency of substitutions related to human adaptation has gradually increased in H3N2 CIVs, and determined four cumulative molecular changes responsible for the increased airborne transmission ability in ferrets. Our results suggested that canine may serve as an intermediate for the adaptation of avian influenza virus to human. Continuous surveillance coordinated with risk assessment for CIVs is necessary.

## Introduction

In the 21st century, newly emerging viruses, such as influenza A H7N9, Ebola virus, Zika virus (ZIKV), and the SARS-CoV-2 virus, have posed serious challenges to health care systems(Jacob et al., 2020, Imai et al., 2017, Yakob and Walker, 2016, Thakur and Ratho, 2022). Those challenges are constant reminders to pay attention to emerging animal-borne zoonotic diseases. Influenza A viruses have a relatively broad host range(Long et al., 2019). When animal-borne viruses with different antigenicity acquire the ability of human-human aerosol transmission, it will become epidemic in the population. The four human pandemic viruses in history underwent avian or swine influenza virus gene reassortment with human influenza virus or acquired human adaptive mutation(Vijaykrishna et al., 2010). Animal-borne viruses adapt to humans through a mammalian intermediate host is an important way to establish infection in humans(Parrish et al., 2015). Swine are considered an intermediate host. For example, the Eurasian avian-origin lineage that originated in European swine in the 1970s gradually accumulated the amino-acid mutations related to human adaptation and increased the infectivity in humans(Mena et al., 2016, Brown, 2013). However, the role of other mammals in ecology is still unclear.

Similar to pigs, the canine respiratory tract contains both types of sialic acid receptors used by influenza viruses (α2,3- and α2,6-linked)(Wasik et al., 2017, Ning et al., 2012). Dogs are susceptible to natural influenza virus infections courtesy of transmission from avian (H3N2 and H5N1), equine (H3N8) or human (pdmH1N1 and H3N2) virus reservoirs(Lin et al., 2012a, Crawford et al., 2005, Song et al., 2008). There have been several events of reassortment of influenza viruses from different host sources in dogs(Lee et al., 2016a, Voorhees et al., 2017), such as ressortment between canine H3N2 and human H1N1 viruses(Song et al., 2012, Moon et al., 2015), and canine H3N2 and swine influenza viruses(Chen et al., 2018). Dogs are important companion animals. Once a new zoonotic disease appears in dogs, there is a high chance to infect humans. However, whether dogs could be as intermediate hosts to produce zoonotic influenza viruses is not sure.

Although dogs have been found to infect with multiple influenza viruses, only equine-origin H3N8 and avian-origin H3N2 viruses have established lineages in dogs(Hayward et al., 2010, Anderson et al., 2012, Zhu et al., 2015, Lyu et al., 2019). Comparing to H3N8 canine influenza virus (CIV), H3N2 CIV has a broader host range, infecting multiple mammalian animals including ferrets, guinea pigs, mice, and cats(Lee et al., 2013, Lyoo et al., 2015, Jeoung et al., 2013, Song et al., 2011). H3N2 CIV was first isolated in 2006 from Guangdong Province in China, and was found to be genetically most closely related to the H3N2 avian influenza viruses prevalent in aquatic birds in South Korea for all eight gene segments (Li et al., 2010, Su et al., 2012). Since then, H3N2 CIV has been prevalent in China(Sun et al., 2013, Yang et al., 2014, Lin et al., 2012b, Wu et al., 2021) and South Korea(Lee et al., 2016b) and has spread to and circulated in the United States since 2015(Voorhees et al., 2018, Martinez-Sobrido et al., 2020, Dalziel et al., 2014). The long-term epidemic of avian-origin H3N2 virus in canine offers the best opportunities to investigate the potential role of dogs in the ecology of influenza A viruses.

Therefore, in the present study, we systematically investigated the evolution of genetic and biological properties of this avian-origin virus during its circulation in dogs. We found that during the adaptation of H3N2 CIVs to dogs, H3N2 CIVs became to recognize the human-like SAα2,6Gal receptor and gradually increased HA acid stability and replication ability in human airway epithelial cells, and had a 100% transmission rate via respiratory droplet in ferret model, which was essential hallmarks of being adapted to infect humans. Our results revealed that dogs might serve as a potential intermediate host for animal influenza viruses to adapt to humans.

## Results

### Continued genetic evolution of avian-origin H3N2 CIVs in dogs

From 2012 to 2019, we collected tracheal swab samples from 4,174 dogs with signs of respiratory disease from animal hospitals and kennels in nine provinces of China (Figure S1). A total of 235 samples (5.63%) were positive for H3N2 infection. The mean positive rates for each year increased from 1.98% in 2012 to 10.85% in 2019 (Figure S2), with a sharp increase after 2016. According to the isolation time and location, 117 representative viruses were selected for full genome sequencing, including 51 strains isolated from 2012-2017 previously uploaded by our laboratory(Lyu et al., 2019). The whole genomes of these viruses were analyzed along with all H3N2 CIVs’ complete genomes publicly available in GenBank and the GISAID database (https://www.gisaid.org/) (n=229), and we constructed the maximum-likelihood phylogenies trees of eight viral gene segments (Figure S3). The inferred trees for all genomic segments exhibited a similar topology. Thus, we grouped them into six clades (clades 0 to 5) according to HA phylogeny, with several viruses consistently clustering together with high posterior probability values (bootstrap values≥70). Clade 0 contains viruses isolated in China from 2006 to 2007, and clades 1 contains isolates exclusively from South Korea from 2007 and 2012. Clade 2 displays viruses from 2009 to 2016 isolated in China and in 2012 collected in Thailand and in 2017 collected in the United States, and clade 3 contains viruses from 2012 and 2013 collected in South Korea. Clade 4 contains strains from 2015 to 2017 isolated in the United States and in 2015 isolated in South Korea, and clade 5 displays isolates from China from 2016 to 2019 and from the United States from 2017 to 2018. Most H3N2 CIVs after 2019 isolated in China further have formed clade 5.1 (Figure 1A). Analysis of sequences of H3N2 CIVs comparing with human influenza viruses and ancestral avian influenza viruses found that, comparing with ancestral avian influenza viruses, H3N2 CIVs which were initially introduced to dogs had possessed several substitutions identical to human influenza viruses with high frequencies (>90%). Noteworthy, the number of human-like amino-acid substitutions has gradually been accumulated during the evolution of H3N2 CIVs in dogs and increased significantly after 2016 (Figure 1B). These results indicated that H3N2 CIVs may increase the adaptability to humans during their evolution in dogs.

**Fig 1.**
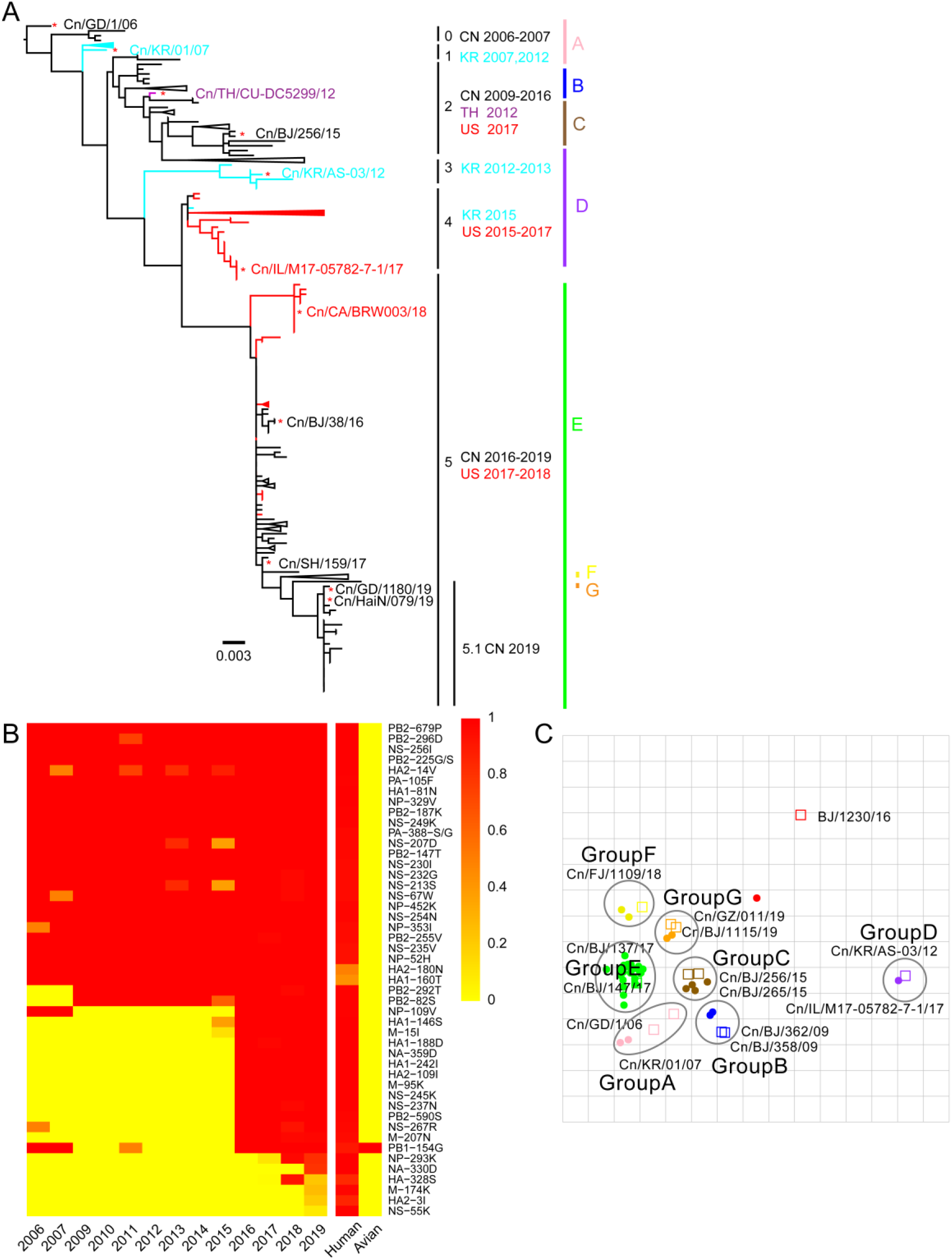
Genetic and antigenic characterization of H3N2 CIVs. (A) Maximum-likelihood phylogenetic tree of hemagglutinin (HA) genomic segment of H3N2 CIVs. The phylogenetic tree of the HA gene was estimated using genetic distances calculated by maximum likelihood under the GTRGAMMA + I model. Viruses labeled with a red “*” were selected for subsequent experiments. Black, red, dark purple, and aqua blue to indicated H3N2 CIVs from China, America, Thailand, and South Korea, respectively. A, B, C, D, E, F and G represent different antigen groups of H3N2 CIVs respectively. A full detailed HA gene tree with the consistent topology is shown in Fig S2 (Scale bar is in units of nucleotide substitutions per site). (B) Prevalence of mammalian adaption markers among H3N2 CIVs. The sequences of H3N2 CIVs available in NCBI were analyzed comparing with avian and human influenza A viruses. Color indicates frequency of indicated substitutions in H3N2 CIVs for each indicated time period. (C) Antigenic map based on the HI assay data. Open squares and filled circles represent the positions of antisera and viruses, respectively. A k-means clustering algorithm identified clusters. Strains belonging to the same antigenic cluster are encircled with an oval. The vertical and horizontal axes both represent antigenic distance. The spacing between grid lines is 1 unit of antigenic distance, corresponding to a two-fold dilution of antiserum in the HI assay. Details of the HI assay data are shown in Table S1.

### Humans lack immunity to H3N2 CIVs

We performed an antigenicity test for representative H3N2 CIVs from different clades. Table S1 and Figure 1C showed that H3N2 CIVs continuously occurred antigenic changes in the worldwide. The cross-reactive titers between different clades were 4-to more-fold lower than homologous reactions. The reaction patterns of clade 0 and clade 1 were similar. They belonged to antigenic group A. Some clade 2 viruses were in antigenic group B, while another clade 2 viruses belonged to antigenic group C. The antigenicity of clade 3 and clade 4 is different from other clades, which belong to antigenic group D. Clade 5 included viruses belonging to antigenic groups E, F, or G. The co-circulation of different antigenic group viruses in recent years increased the difficulty of preventing and controlling canine influenza viruses. Additionally, we found that all of H3N2 CIVs were not recognized by antisera of H3N2 human seasonal influenza virus.

To further investigate whether humans have exiting immune protection against H3N2 CIVs. We tested sera collected from children (< 15 y old, n = 100), adults (25– 53 y old, n = 100), and elderly adults (≥ 60 y old, n =100) against four viruses (BJ/1230/16, human seasonal H3N2 influenza; Cn/BJ/38/16, group C; Cn/FJ/1109/18, group D and Cn/GZ/011/19, group E) for both HI and microneutralization (MNT) antibodies, as described previously(Potter and Oxford, 1979, Rowe et al., 1999). We found that 15.0%, 1.0%, 1.0%, and 2.0% of the children; 8.0%, 0.0%, 1.0%, and 1.0% of the adults; and 5.0%, 0.0%, 0.0%, and 1.0% of the elderly adults had HI antibody titers of ≥ 40 to BJ/1230/16, Cn/BJ/38/16, Cn/FJ/1109/18, and Cn/GZ/011/19, respectively (Table S2). In addition, 14.0%, 1.0%, 1.0%, and 2.0% of the children had MNT antibody titers of ≥ 40, and 6.0%, 0%, 0%, and 0% of the adults and 4.0%, 0%, 0%, and 0% of the elderly adults had MNT antibody titers of ≥ 80 to BJ/1230/16, Cn/BJ/38/16, Cn/FJ/1109/18, and Cn/GZ/011/19, respectively (Table S2). These results indicated that preexisting immunity derived from the present human seasonal influenza viruses cannot provide protection against H3N2 CIVs.

### H3N2 CIVs obtained human-type receptor-binding properties and their acid stability increased stepwise

Our genetic analysis found that humanized adaptive mutations increased significantly along with the prevalence of H3N2 CIVs in dogs, while humans lack preexisting immunity to the H3N2 CIVs indicating that H3N2 CIVs might spread in human populations once they adapted to humans. Therefore, we further evaluated the potential threat of H3N2 CIVs for public health. The binding preference of HA for the host SAα2 6Gal receptor and low activation pH are critical determinants for cross-species transmission of influenza virus to humans (Connor et al., 1994, Matrosovich et al., 2000). Therefore, we examined the receptor-binding preference of 11 H3N2 canine influenza viruses isolated from 2006 to 2019 (Figure S4). We noticed that compared with the H3N2 avian influenza virus Dk/KR/JS53/04 and H3N2 CIVs from clades 0, 1, 2, and 3, which only recognized α-2,3-linked sialosides (Figure 2A), H3N2 CIVs belonged to clade 4, 5 and 5.1, represented by Cn/US/M17/17, Cn/SH/159/17 and Cn/Hain/079/19 showed dual binding specificity to both α-2,3- and α-2,6-linked sialosides. Assay of haemagglutinin (HA) acid stability showed that the HA acid stability of clade 5 viruses (activation pH 5.3) was higher than clade 0, 1, 2, 3, and 4 viruses (activation pH 5.4) (Figure 2B and Figure S5). Compared with the H3N2 CIVs that circulated before 2019 (activation pH 5.3), the viruses circulating after 2019 belonging to clade 5.1 had a higher HA acid stability (activation pH 5.2) and was identical to human H3N2 virus HA fusion pH.

**Fig 2.**
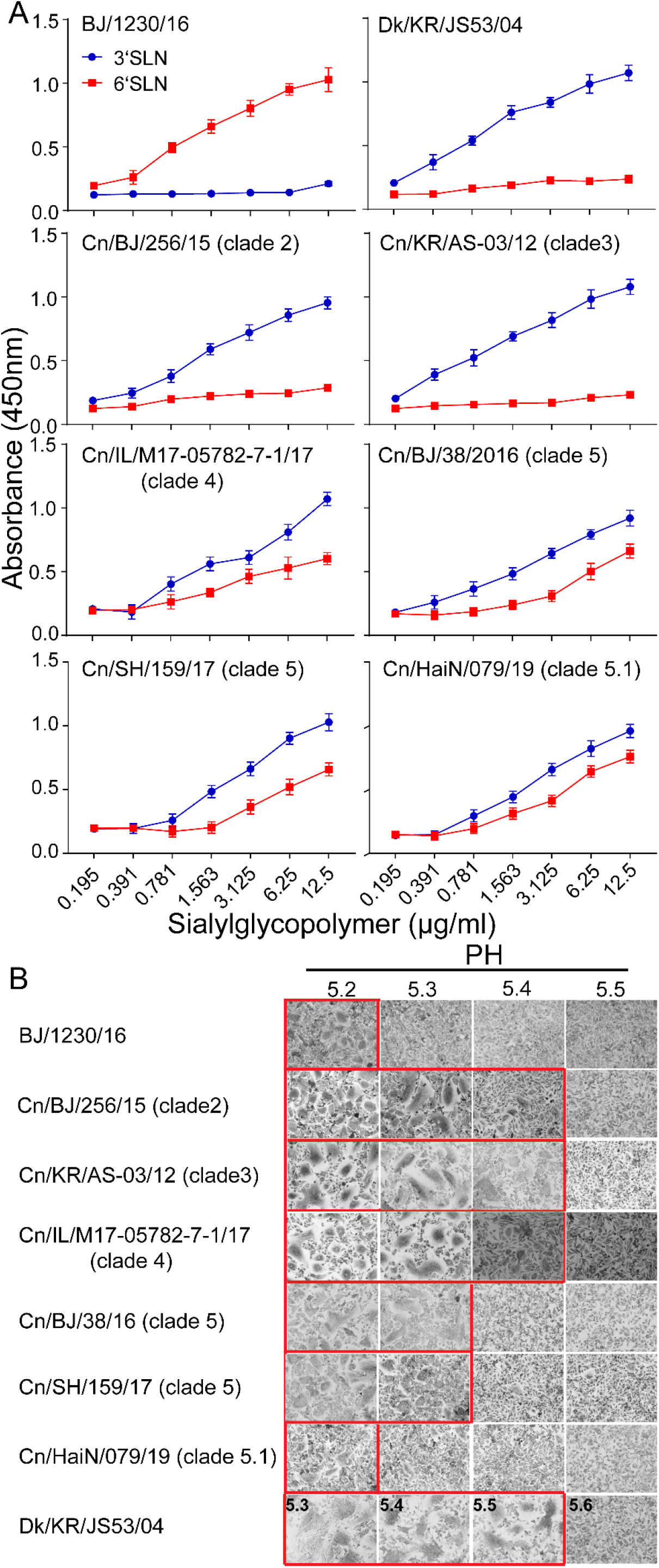
Binding specificities toward α-2, 3- or α-2, 6-linked sialic acid receptors and HA acid stability. (A) Characterization of the receptor-binding properties of H3N2 CIVs. The direct binding of the virus to sialylglycopolymers containing either 2,3-linked (blue) or 2,6-linked (red) sialic acids was tested (n=3 biological replicates and n=3 technical replicates). (B) HA activation pH measured by syncytia assay. Representative fields of cells infected with the indicated viruses and exposed to pH 5.2, 5.3, 5.4, 5.5, or 5.6 are shown. Images were taken at ×10 magnification. The experiments were repeated three times, with similar results.

The replication efficiency of CIVs exhibiting different receptor binding specificity and HA acid stability were evaluated *in vitro*. In A549 cells, the titers of clade 5 and 5.1 viruses were significantly higher (up to nearly 100-fold higher) than other clades over 12 h to 72 h (*P* < 0.01) (Figure 3A), and clade 5.1 viruses showed comparable virus outputs with the human seasonal H3N2 virus at each time point. Infection of NHBE cells with H3N2 CIVs produced similar progeny results. The titers of clade 5 and 5.1 viruses in NHBE cells were significantly (up to nearly 100-fold higher) higher than viruses from other clades between 24 hpi and 72 hpi (*P* < 0.01), and clade 5.1 viruses and human seasonal H3N2 virus also replicated to similar levels at each time point (Figure 3B). Collectively, H3N2 CIVs obtained human-type receptor binding properties, and their HA acid stability and replication ability in human cells increased stepwise during their circulation in dog population.

**Fig 3.**
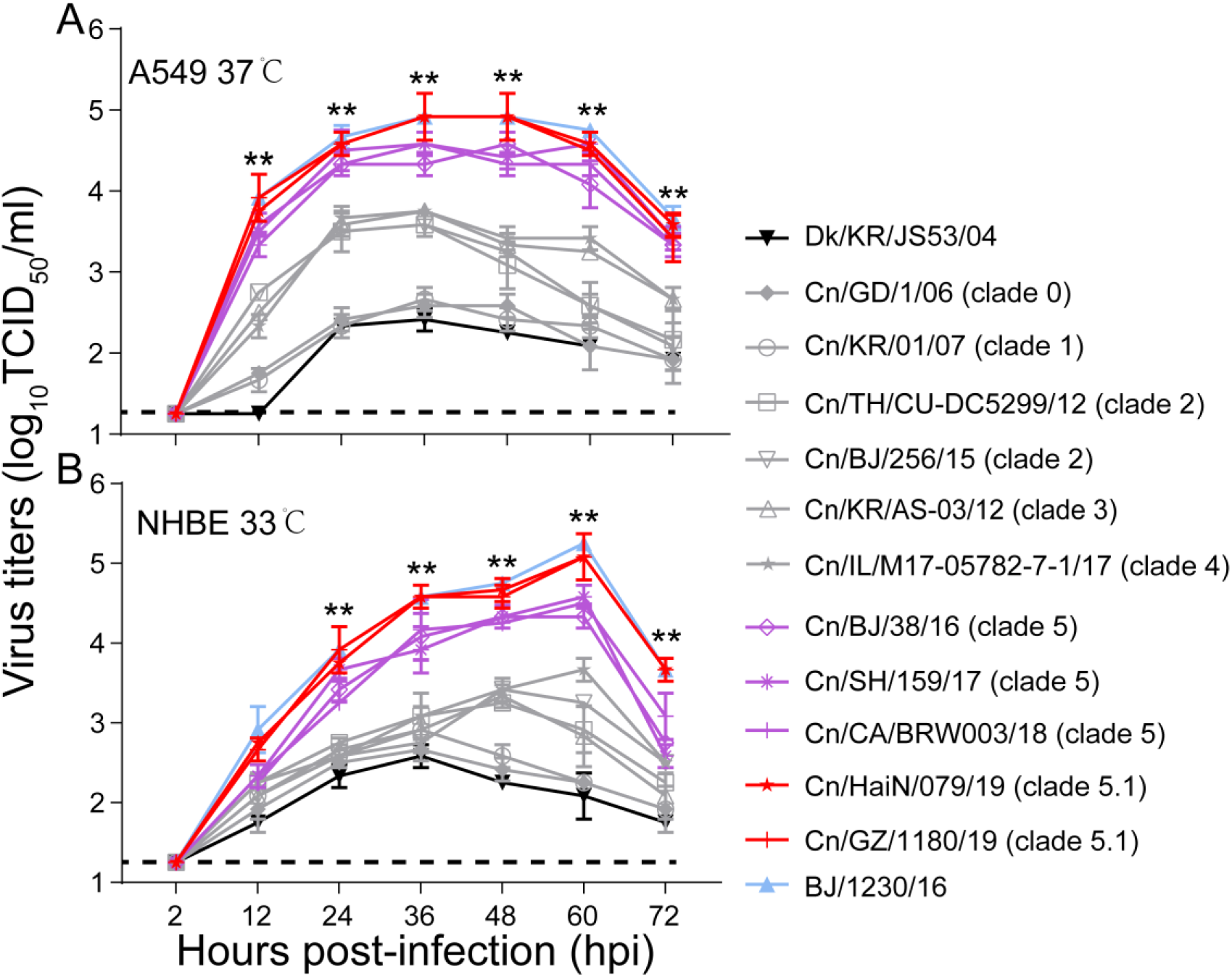
Viral growth properties in A549 (A) and NHBE (B) cells. Cells were infected with indicated viruses at MOI of 0.01 and incubated at 37°C or 33°C. Supernatant s were harvested at the indicated time points, and the virus titers were determined in MDCK cells. Values are expressed as means ± standard deviations (SD) (n=3 biological replicates and n=3 technical replicates). **, P < 0.01, the titers of BJ/1230/16 (human seasonal H3N2 virus), clade 5 or 5.1 viruses were significantly higher than other viruses. The dashed black lines indicate the lower limit of detection.

### Improved replication and transmissibility of H3N2 CIVs in dogs

To evaluate the infectivity and transmission ability of H3N2 CIVs in dogs, we inoculated intranasally six dogs with 10^6^ TCID_50_ of each virus strain. Twenty-four hours later, three inoculated dogs for each virus strain were individually paired and cohoused with a direct-contact dog. We found that H3N2 avian influenza virus (Dk/KR/JS53/04) and H3N2 CIVs (from clades 0, 1, 2, 3, and 4) caused only mild clinical signs with mean clinical scores ranging from 0.5 to 2.5 and a mean body temperature ranging from 37.7°C to 39.7°C (Figure 4A and 4B). However, infection with H3N2 CIVs from clade 5 and 5.1 resulted in more severe clinical symptoms such as pyrexia, sneezing, wheezing, and coughing, with a higher mean clinical score ranging from 2.8 to 3.3 and a higher mean body temperature ranging from 39.9°C to 40.2°C. H3N2 avian influenza virus cannot be detected in dogs at 4 dpi, while H3N2 CIVs from clade 5 and 5.1 can replicate efficiently in both the upper (nasal turbinate and trachea) and lower (lung) respiratory tracts and tonsils of dogs which were significantly higher than those of other clades viruses (*P* < 0.01) (Figure 4C). Furthermore, the virus identification of nasal swabs showed that H3N2 CIVs was efficiently transmitted to all naive dogs by direct contact (Figure 4D and Figure S6). In contrast, the H3N2 avian influenza virus cannot transmit between dogs. H3N2 CIVs (from clade 0, 1, 2, 3, and 4) could be detected in two of three naïve animals or all contact animals at 4 dpi or 6 dpi, and seroconversion was detected in 2/3 contacts (1:80 to 1:160) or all contacts (1:160 to 1:320). Noteworthy, clade 5 and 5.1 viruses, represented by Cn/SH/159/17 and Cn/HaiN/079/19, were transmitted to all three contact animals at 2 dpi with seroconversion in all contacts (1:320 to 1:640), which was earlier than clade 0, 1, 2, 3 and 4 viral transmission to the naïve animals. Furthermore, clade 5 and 5.1 viruses also replicated more efficiently in all donors than other clades viruses (*P* < 0.01). Thus, the replication and transmissibility of H3N2 CIVs gradually increased in dogs.

**Fig 4.**
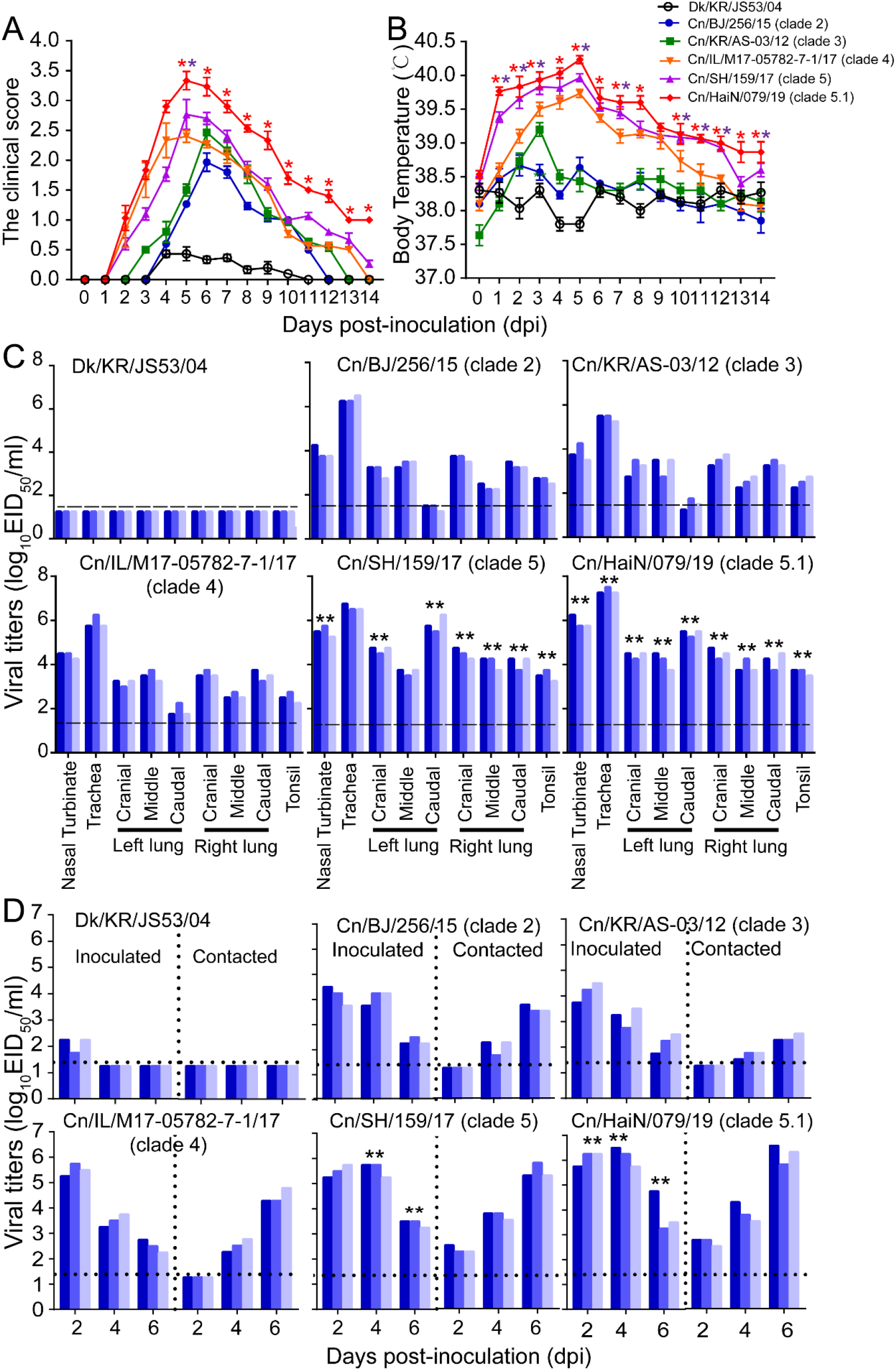
Infectivity and transmissibility of H3N2 CIVs in dogs. (A) Clinical symptoms score of dogs infected with H3N2 CIVs. (B) Body temperatures of dogs infected with H3N2 CIVs. (C) Virus replication in the indicated organ. Three dogs were infected intranasally with 10^6^ EID_50_ of each virus and euthanized at 4 dpi for virus enumeration. Viral titers in the nasal turbinates, tracheas, lungs, and tonsils were determined by the infection of eggs. Each color bar represents the virus titer of an individual animal. (D) Direct contact transmission of H3N2 CIVs in dogs. Statistical significance of clade 5 or clade 5.1 viruses relative to other viruses in the inoculated animals were assessed using two-way ANOVA (*, *P* < 0.05; **, *P* < 0.01).

### H3N2 CIVs acquired efficient aerosol transmissibility in the ferret model after 2016

To further evaluate the potential risk of the H3N2 CIVs to the public health, we examined the replication and transmission of H3N2 viruses in a ferret model. A group of six ferrets was inoculated intranasally with 10^6^ TCID_50_ of each virus strain. Twenty-four hours later, three inoculated ferrets for each virus strain were individually paired. An uninfected animal was housed in a wire-frame cage adjacent to the infected ferret to assess aerosol spread. Virus detection in nasal washes of the aerosol-spread animal showed that the H3N2 avian influenza virus (Dk/KR/JS53/04) and H3N2 CIVs (from clade 0, 1, 2, 3 and 4) did not transmit to ferrets by respiratory droplets (Figure 5). Clade 5 viruses, represented by Cn/BJ/38/16, Cn/SH/159/17, and Cn/US/BRW003/18, transmitted to all naïve three ferrets via respiratory droplets at 6 dpi, and seroconversion was detected in all aerosols (1:160 to 1:320). More importantly, clade 5.1 viruses, represented by Cn/HaiN/079/19 and Cn/GZ/1180/19, were able to transmit to all three naïve ferrets as early as 4 dpi with seroconversion in 3/3 aerosols (1:320 to 1:640). In addition, clade 5 and 5.1 viruses also replicated more efficiently in all donors than other clades viruses. Clade 5.1 viruses showed comparable virus outputs and transmissibility with the human seasonal H3N2 virus in ferret model. Collectedly, H3N2 CIVs obtained aerosol transmissibility during their evolution in dogs.

**Fig 5.**
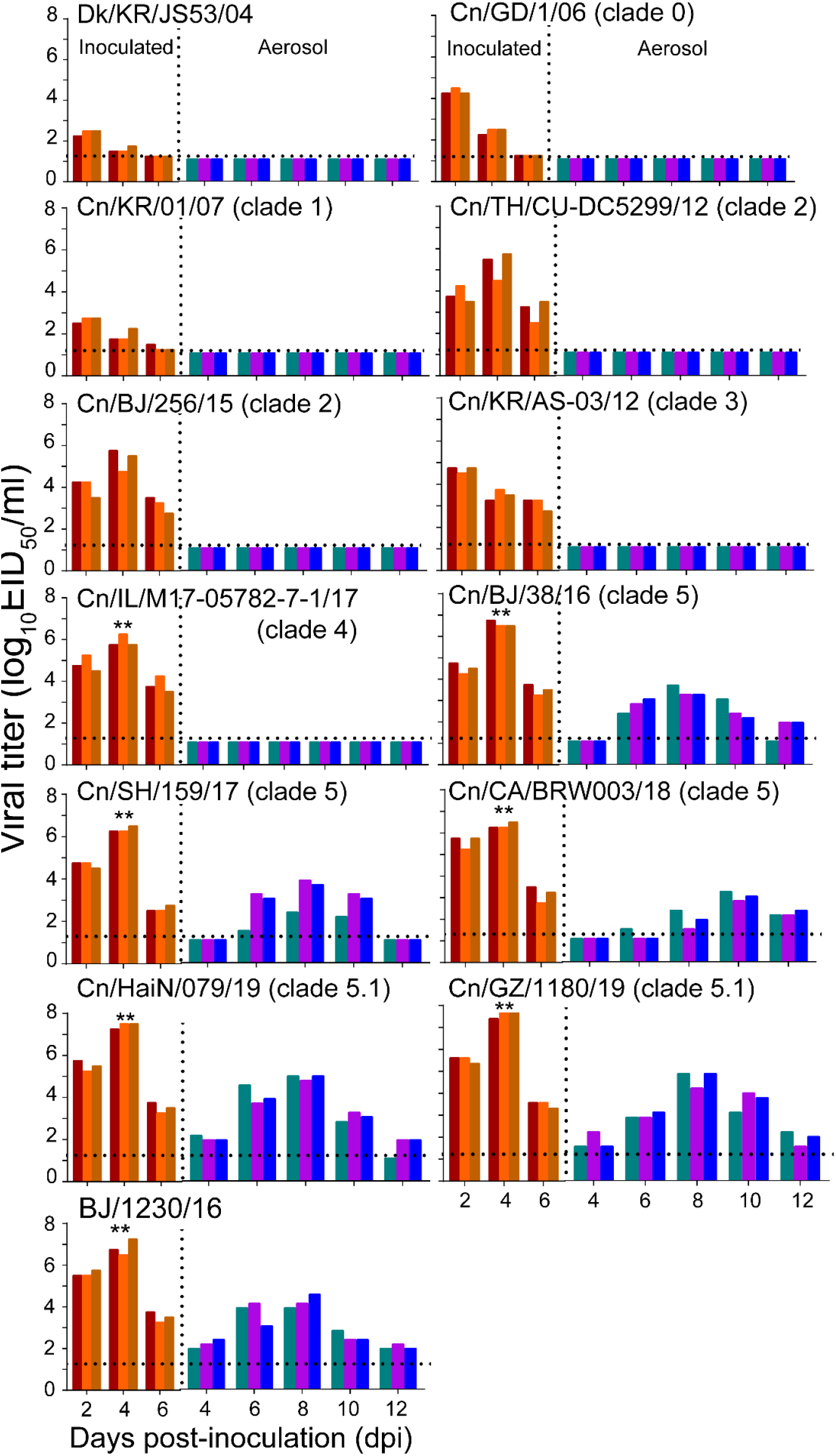
Respiratory droplet transmission of H3N2 CIVs in ferrets. Groups of three ferrets were infected intranasally with 10^6^ EID_50_ of indicated viruses and then housed separately in solid stainless-steel cages within an isolator. The next day, three uninfected respiratory droplet contact animals were individually housed in a wire-frame cage adjacent to the infected ferret. Nasal washes were collected every other day from all animals for virus shedding detection from day 2 of the initial infection. Each color bar represents the virus titer of an individual animal. No data are displayed when the virus was not detected from all of the groups. Dashed lines indicate the lower limit of virus detection. Statistical significance of human influenza virus (BJ/1230/16), clade 5.1 or clade 5 viruses relative to other viruses in the inoculated animals were assessed using two-way ANOVA (**, *P* < 0.01).

### Molecular determinants associated with efficient transmission reside in the HA and PB1 genes

Since recent strains have increased replication and transmissibility in dogs and have gradually acquired 100% aerosol transmission ability in the ferret model, we further determined the molecular mechanisms responsible for the enhanced replication and transmission of H3N2 CIVs in mammals. We used Cn/BJ/256/15 (clade 2) as the backbone to generate reassortant viruses by single replacing all genes from the Cn/HaiN/079/19 (clade 5.1) virus and tested their replication and transmissibility among dogs and ferrets. We found that PB1 and HA genes significantly enhanced the replication and transmission of single-gene reassortant virus both in dogs and ferrets (Table S3 and S4), and seroconversion was detected in all contact dogs (1:160 to 1:320) and 2/3 aerosol ferrets (1:80 to 1:160). Next, we identified 15 conserved amino acid variations in the HA protein and PB1 protein of H3N2 CIVs that emerged in 2016-2019 (Figure S7). Among them, HA-146S, 188D, and PB1-154G were also highly enriched (>90%) among human H3N2 influenza A viruses isolated from 2006 to 2019 (Figure 6A). In addition, HA-16S was detected at a high frequency (>81%) among the H3N2 CIVs isolated after 2019, while all of H3N2 CIVs before 2018 possessed HA-G16 (Figure 6A). Next, we evaluated the effect of these substitutions on receptor binding property assays and acid stability and thermal stability experiments. We found that introduction of the HA-G146S enhanced binding for α-2,6-linked sialosides of Cn/BJ/256/15 (Figure 6B). The introduction of the HA-G16S and HA-N188D increased the HA acid and temperature stability of Cn/BJ/256/15 (Figure 6C and 6D). The introduction of the PB1-D154G increased the polymerase activity of Cn/BJ/256/15 (Figure 6E).

**Fig 6.**
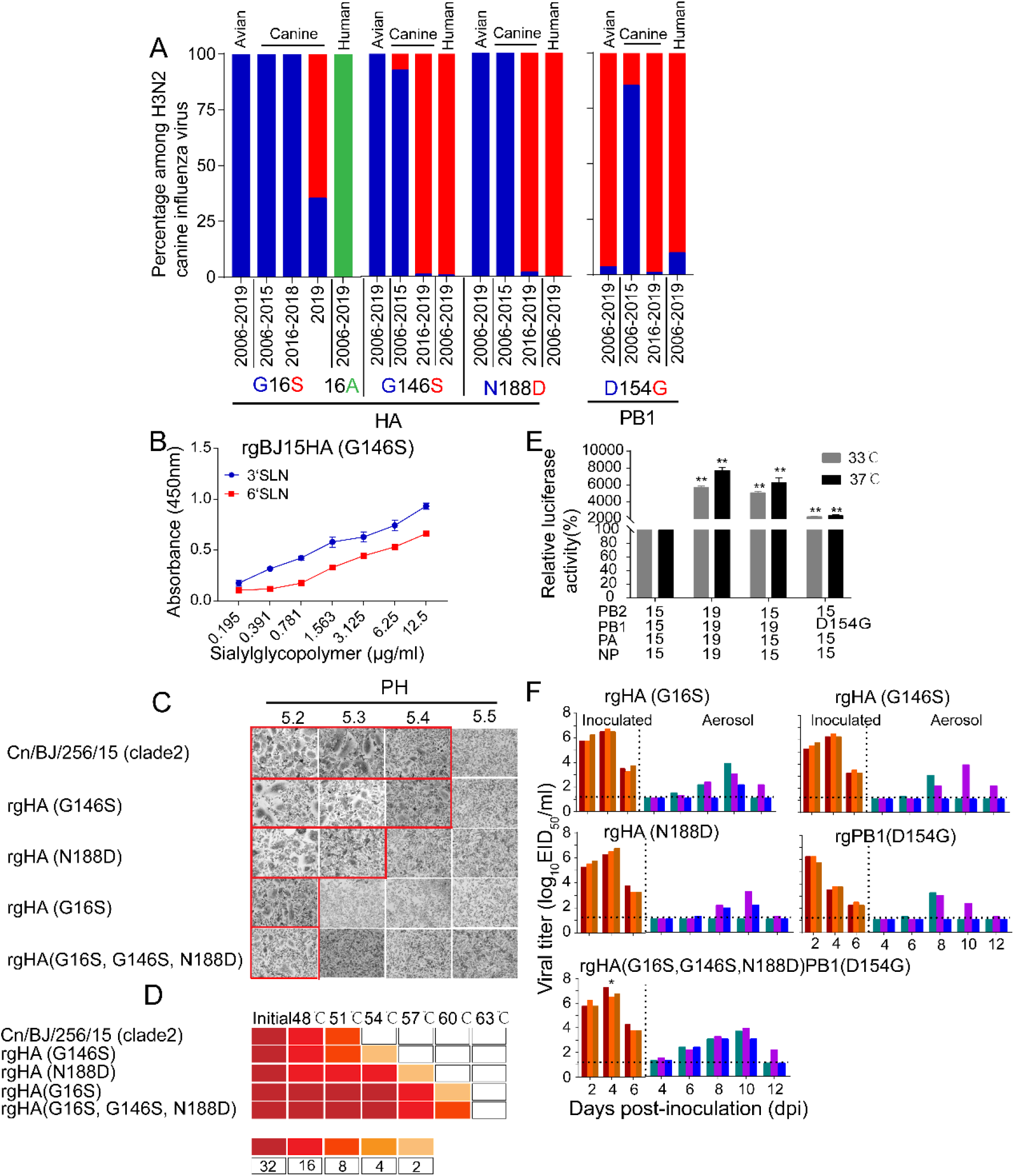
The HA-G16S, G146S, N188D, and PB1-D154G mutations in H3N2 CIVs were the minimal molecular change required to facilitate the efficient aerosol transmissibility of the non-transmissible clade 2 virus. (A) Detection frequency of G/S at HA residue 16 and 146, N/D at HA1 residue 188 (n= 437), and D/G at PB1 residue 154 (n=437) in avian, canine, and human influenza viruses. Amino acid residues are colored in blue, red, and green, respectively. (B) Identification of mutations that confer binding to human-type receptors.(C) Epresentative fields of vero cells expressing the indicated HAs and exposed to pH 5.2, 5.3, 5.4, or 5.5 are shown. Images were taken at ×10 magnification. The experiments were repeated three times, with similar results. (D) HA protein stability as measured by the ability of viruses to agglutinate CRBCs after incubation at indicated temperatures for 40 min. Colors indicate the hemagglutination titers upon treatment at various temperatures for 40 min, as shown in the legend. The experiments were repeated three times with similar results. (E) The effects of PB1 from 19(Cn/HaiN/079/19) and amino-acid substitutions at PB1 residue 154 on the viral polymerase activity were determined using a minigenome assay in 293T cells. Values shown are the mean ± SD of the three independent experiments and are standardized to those of 15(Cn/BJ/256/15) measured at 37°C(100%) and 33°C(100%). Statistical significance was assessed using two-way ANOVA (**, *P* < 0.01). (F) Aerosol transmissibility of the mutation viruses in ferrets. Statistical significance of rgHA(G16S, G146S, N188D)PB1(D154G) viruses relative to other other single substitution viruses were assessed using two-way ANOVA (*, *P* < 0.05).

We then focused on these four mutations and evaluated the replication and respiratory droplet transmission ability of rgHA-(G16S), rgHA(G146S), rgHA(N188D), rgPB1(D154G), and rgHA(G16S, G146S, N188D)PB1(D154G) viruses in ferrets. We found that rgHA(G146S), rgHA(N188D), and rgPB1(D154G) viruses were transmitted to two of the three ferrets with seroconversion in 2/3 aerosols (1:160 to 1:320), and the rgHA-(G16S) and rgHA(G16S, G146S, N188D)PB1(D154G) viruses were transmitted to all three ferrets with seroconversion in 3/3 aerosols (1:320 to 1:640) (Figure 6F). In addition, the viral titers in the nasal washes from the rgHA(G16S, G146S, N188D)PB1(D154G)-inoculated animals were higher than that of the other single substitution viruses (*P* < 0.05) and were similar to transmissible wild-type virus (Cn/HaiN/079/19). These results indicated that HA-G16S, G146S, N188D, and PB1-D154G are crucial for the replication and efficient transmissibility of H3N2 CIVs in ferrets.

## Discussion

By evaluating the biological characteristics of avian-origin H3N2 CIVs isolated in dogs from different years, we found that the effective infection and transmission in dogs was not an intrinsic property possessed by H3N2 CIVs but through stepwise adaptation. Of note, threats to human health increased during H3N2 CIVs adaptation to dogs. Specifically, we observed changes in receptor binding specificity from recognizing only α-2,3-linked sialosides to recognizing both α-2,3- and α-2,6-linked sialosides and gradually increased HA acid stability and replication in human airway epithelial cells and ferrets, and can transmit by aerosol among ferrets. In addition, humans lack immunity to the H3N2 CIVs, and increased isolation of H3N2 CIVs in the dog population increased the chance of transmission to humans. Therefore, we suggested that dogs was a potential intermediate host of avian influenza viruses to adapt to humans.

We found that the H3N2 CIVs formed different clades after entering the dog population, and these clades were related to the time of virus isolation. Early H3N2 CIVs (clade 0 and 1) do not transmit efficiently in dogs. After a particular opportunity, the avian H3N2 subtype influenza virus enters the dog and gradually adapts to the dog. The H3N2 CIVs isolated after 2012 can reach 100% transmission in dogs. Clade 5 viruses isolated after 2016 were transmitted to all three contact animals within 2 dpi. The transmission time interval is shortened. Moreover, the clade 5.1 replication ability generated after 2019 has been further improved. Besides, the antigenicity of H3N2 CIVs have continuously evolved into groups A, B, C, D, and E. The gradual improvement of infection and transmission ability and the continuous evolution of antigenicity promoted the prevalent of H3N2 CIV in dogs.

The high isolation rate of the H3N2 CIVs in companion animal dogs increased the opportunity of viral cross-species transmission to humans. We found that the number and frequency of substitutions identical to human influenza viruses increased in H3N2 CIVs during the adaptation process of dogs, and the homology sites increased significantly after 2016. Therefore, we evaluated the potential threat of H3N2 CIVs, and found that due to the mutation of HA-G146S around the receptor-binding domain after 2016 from recognition of only α-2,3-linked sialosides (clade 0, 1, 2, and 3 viruses) to both α-2,3- and α-2,6-linked sialosides (clade 4, 5, and 5.1). Besides, the HA-G16S and HA-N188D mutation resulted in enhanced acid and thermostability, and the PB1-D154G mutation resulted in increased polymerase activity, which together resulted in the acquisition of aerosol-transmitting properties in ferrets. Noteworthy, the proportion of HA-G146S, HA-N188D, and PB1-D154G are all higher than 90% in human influenza viruses. Therefore, during the gradual adaptation of CIVs in dogs, clade 5 CIVs with high adaptability to mammals was selected. Additionally, we found that clade 5 CIVs further adapted after 2019. HA-G16S mutation which improved acid stability further improved replication level and shortened transmission time in ferrets.

Taken together, our results suggests that after the avian-origin H3N2 influenza viruses were introduced to dogs, they adapted to canines and increased the threat to public health. Controlling the prevailing H3N2 CIVs in dogs and continuous monitoring their biological characteristics should be implemented.

## Materials and methods

### Ethics statements and facility

The present study was carried out according to the Guide for the Care and Use of Laboratory Animals of the Ministry of Science and Technology of the People’s Republic of China. The protocols for the animal studies were approved by the Committee on the Ethics of Laboratory Animals of China Agricultural University (approval SKLAB-B-2010-003).

### Viruses and cells

Nasopharyngeal swabs were collected from dogs with signs of respiratory disease in animal hospitals or kennels in nine provinces of China (Liaoning, Beijing, Tianjin, Nanjing, Shanghai, Shanxi, Guangzhou, Fujian, and Hainan) between 2017 and 2019. Nasopharyngeal swabs were placed in 1.0 ml of transmission medium [50% (vol/vol) glycerol in PBS] containing antibiotics, as previously described (Zhang et al., 2008). We amplified the matrix gene by real-time reverse transcription (RT) PCR using the Influenza A Virus V8 Rapid Real-Time RT-PCR Detection Kit (Beijing Anheal Laboratories Co. Ltd., http://anheal.company.weiku.com), and isolated and identified virus isolates using methods described previously (Sun et al., 2013). The Chinese H3N2 CIVs we isolated before 2017 and human seasonal H3N2 virus A/Beijing/1230/2016 (BJ/1230/16) were described previously (Lyu et al., 2019, Sun et al., 2013, Sun et al., 2021).

HEK293T cells, human lung carcinoma cells (A549), Vero cells, and Madin-Darby canine kidney (MDCK) cells were cultured in Dulbecco’s modified Eagle’s medium (DMEM Gibco) supplemented with 10% fetal bovine serum (FBS; Gibco), 100 U/ml of penicillin, and 100 μg/ml of streptomycin. NHBE cells (Lonza, Allendale, NJ, USA) were cultured in bronchial epithelial cell growth medium (Lonza) at the air-liquid interface, as previously described (Hudy et al., 2010, Matrosovich et al., 2004). All cells were maintained in a humidified incubator containing 5% CO_2_ at 37°C (HEK293T, A549, Vero cells, and MDCK cells) or 33°C (NHBE cells).

### Genetic and phylogenetic analyses

Viral gene amplification and sequencing were carried out as previously described (Sun et al., 2010). All previously published gene sequences from the H3N2 canine influenza virus were collected from the NCBI and the Global Initiative on Sharing Avian Influenza Data (https://www.gisaid.org). Phylogenetic analyses were performed on regions of the alignments containing the fewest gaps across sequences. These regions consisted of the following intervals: PB2, 82–2,061; PB1, 83–2,167; PA, 82–2,068; HA, 34–1,656; NP, 13–1,446; NA, 36–1,377; MP, 67–924; and NS, 28–838. RAxML was used to construct maximum likelihood phylogenies for each segment (Stamatakis, 2014), via CIPRES Science Gateway(Miller et al., 2010); 1,000 bootstrap replicates were run, and GTRGAMMA + I was used as the nucleotide substitution model.

### Generation of recombinant viruses by reverse genetics

The genome sequence of avian influenza virus A/duck/Korea/JS53/2004 (Dk/KR/JS53/04) and H3N2 CIVs A/canine/Guangdong/1/2006 (Cn/GD/1/06), A/canine/Korea/01/2007 (Cn/KR/01/07), A/canine/Thailand/CU-DC5299/2012 (Cn/TH/CU-DC5299/12), A/canine/Korea/AS-03/2012 (Cn/KR/AS-03/12), A/canine/lllinois/M17-05782-7-1/2017 (Cn/IL/M17-05782-7-1/17) and A/canine/California/BRW003/2018 (Cn/CA/BRW003/18) were downloaded from the NCBI influenza virus database and synthesized by Sangon Biotech Company (Shanghai, China). The eight gene segments of each virus were amplified by RT-PCR and cloned into the dual-promoter plasmid pHW2000, and then these viruses were generated using reverse genetics as previously described (Sun et al., 2011).

All eight gene segments from H3N2 CIVs A/canine/Beijing/0118-256/15(Cn/BJ/256/15, BJ15) and A/canine/Hainan/079/2019 (Cn/HaiN/079/19, HaiN19) were amplified by RT-PCR and then cloned into the dual-promoter plasmid pHW2000. Reverse-genetics systems for BJ15 and HaiN19 were then established, and single-gene reassortant viruses were rescued by reverse genetics. Mutations were introduced into the PB1and HA genes using a QuikChange site-directed mutagenesis kit (Agilent) in accordance with the manufacturer’s instructions. PCR primer sequences are available upon request. All constructs were sequenced to ensure the absence of unwanted mutations. The HA- and PB1-mutant viruses were generated using reverse genetics. Rescued viruses were detected using hemagglutination assays. Viruses were purified by sucrose density gradient centrifugation. Viral RNA was extracted and analyzed by RT-PCR, and each viral segment was sequenced.

### Antigenic analyses

Antigenic characteristics of H3N2 viruses were compared using an HI assay with ferret antisera raised against representative viruses. Ferret antisera raised against BJ/1230/16, Cn/BJ/358/09 (group A), Cn/BJ/362/09 (group A), Cn/BJ/256/15 (group B), Cn/BJ/265/15 (group B), Cn/BJ/137/17 (group C), Cn/BJ/147/17 (group C), Cn/FJ/1109/18 (group D), Cn/GZ/011/19 (group E), and Cn/BJ/1115/19 (group E) viruses were produced in our laboratory. All HI assays were performed following WHO guidelines and in duplicate. Antigenic properties were analyzed using the antigenic cartography methods described previously(Smith et al., 2004). The antigen cartography of the viruses was constructed using Antigenic Cartography software (http://www.antigenic-cartography.org/). Clusters were identified in the antigenic map by a k-means clustering algorithm using average weighting and k = 3. The human sera were selected from children, adults, and elderly adults who visited the hospital but did not have a fever. Informed consent to use the sera for influenza antibody detection was obtained. Sera were treated with *Vibrio cholerae* receptor-destroying enzyme (Denka-Seiken) before being tested for the presence of HI antibody with 1% CRBC. The minimum cut-off value for the HI assay was 10. MNT assays were performed as reported previously(Rowe et al., 1999, Hancock et al., 2009).

### Receptor-binding assays

α-2,6 glycans (6′ SLN: Neu5Acα2-6Galβ1-4GlcNAc β-SpNH-LC-LC-biotin) and α-2,3 glycans (3′ SLN: Neu5Acα2-3Galβ1-4GlcNAcβ-SpNH-LC-LC-biotin) were kindly provided by the Consortium for Functional Glycomics (Scripps Research Institute, Department of Molecular Biology, La Jolla, CA). Receptor-binding specificity was determined by a solid-phase direct binding assay as previously described(Chandrasekaran et al., 2008). Briefly, serial dilutions (0.195 μg/ml, 0.391 μg/ml, 0.781 μg/ml, 1.563 μg/ml, 3.125 μg/ml, 6.25 μg/ml, and 12.5 μg/ml) of biotinylated glycans 3′SLN and 6 ′SLN were prepared in PBS. After, 100 μl was added to the wells of 96-well microtiter plates and allowed to attach overnight at 4°C. The plates were then irradiated with 254 nm ultraviolet light for 2 min. After removing the glycopolymer solution, the plates were blocked with 0.1 ml of PBS containing 2% bovine serum albumin (BSA) at room temperature for 1 h. After washing with ice-cold PBS containing 0.1% Tween 20 (PBST), the plates were incubated in a solution containing influenza virus (64 HA units in PBST) at 4°C for 12 h. After washing with PBST, chicken antisera against Dk/KR/JS53/04 (avian H3N2), Cn/BJ/137/17 (canine H3N2), or BJ/1230/16 (human H3N2) virus was added to each well, and the plates were incubated at 4°C for 2 h. The wells were then washed with ice-cold PBST and incubated with HRP-linked goat anti-chicken antibody (Sigma-Aldrich) for 2 h at 4°C. After washing with ice-cold PBST, the plates were incubated with O-phenylenediamine in PBS containing 0.01% H_2_O_2_ for 10 minutes at room temperature. The reaction was stopped with 0.05 ml of 1 M H_2_SO_4_, and the absorbance was determined at 450 nm.

### Viral growth kinetics in cells

The TCID_50_ was determined in MDCK cells by inoculation of 10-fold serially diluted viruses at 37°C for 48 h. The TCID_50_ value was calculated by the Reed-Muench method. Multistep replication kinetics were determined using A549 and NHBE cells. A549 cells were infected with viruses at an MOI of 0.01, overlaid with serum-free DMEM containing1g/ml tosylsulfonyl phenylalanyl chloromethyl ketone (TPCK)-trypsin (Sigma-Aldrich), and incubated at 37°C. NHBE cells were infected with viruses at an MOI of 0.01 and cultured in B-ALI growth medium (Lonza) at 33°C. The supernatants were sampled at 2, 12, 24, 36, 48, 60, and 72 hpi and titrated by inoculating MDCK cells in 96-well plates. Three independent experiments were performed.

### Dog pathogenesis and transmission experiments

We used 10-week-old female beagles (Beijing Marshall Biotechnology Co., Ltd) seronegative for currently circulating influenza viruses as study subjects. The dogs were anesthetized with ketamine (20 mg/kg), xylazine (1 mg/kg), and infected intranasally with 10^6.0^ EID_50_ of test virus in a 2-ml volume. The animals were subsequently euthanized at 4 dpi, and nasal turbinate, trachea, lung, and tonsil samples were collected for virus titration. Lung tissues were also used for pathological examination.

For transmission studies, groups of three female beagles housed in a cage placed inside an isolator were inoculated intranasally with 10^6.0^ EID_50_ of test virus. Twenty-four hours later, the three inoculated animals were individually paired by co-housing with a direct-contact dog. Nasal swabs were collected at 2-day intervals from 2 dpi to 14 dpi. Nasal secretion swabs were taken and placed in a 1.0ml transmission medium [50% (vol/vol) glycerol in PBS] containing antibiotics. Viruses in the nasal secretion swabs were titrated in eggs. Sera were collected from contacted animals at 21 dpi. Seroconversion was analyzed by HI assay. Clinical signs and temperature were recorded daily for all inoculated dogs.

### Ferret pathogenesis and transmission experiments

Six-month-old female Angora ferrets (Wuxi Sangosho Biotechnology Co., Ltd, Angora) seronegative for currently circulating influenza viruses were used. In the transmission experiment, groups of three animals were each anesthetized with ketamine (20mg/kg) and xylazine (1mg/kg), and infected intranasally with 10^6.0^ EID_50_ of test virus in a 500-μl volume (250μl per nostril). Twenty-four hours later, the three inoculated animals were paired with naïve ferrets, housed in a wire-frame cage adjacent to the infected ferrets. The infected and respiratory droplet ferrets were 5 cm apart. Nasal washes were collected at 2-day intervals from 2dpi to 14dpi and titrated for viruses using eggs to monitor virus shedding. Sera were collected from respiratory droplet animals at 21 dpi. Seroconversion was analyzed by HI assay. The ambient conditions for these studies were 20 to 22°C and 30% to 40% relative humidity. The airflow in the isolator was horizontal at a speed of 0.1 m/s; the airflow direction was from the inoculated animals toward the exposed animals.

### HA thermostability

Purified viruses were diluted in PBS to 32 HA units and dispensed by 120μl into 0.2-ml, thin-walled PCR tubes (USA Scientific, Ocala, FL). Tubes were placed into a Gradient Veriti 96-well thermal cycler (catalog number 9902; Life Technologies, Camarillo, CA). The temperature range was set at 48°C to 63.0°C. Tubes were heated for 40 min and then transferred to ice 5min. Control samples containing 120μl of virus were incubated for 40 min at 0°C. The virus content in each sample was determined by an HA assay using a 1% suspension of chicken erythrocytes. Each virus sample was analyzed three times for thermostability.

### HA acid stability

HA acid stability was measured using syncytia assays. Vero cells in 24-well plates were infected at 3 PFU/cell MOI in the syncytia assay. At 16 hpi, infected cells were treated with DMEM supplemented with 5 mg/ml TPCK-treated trypsin for 15 min. Infected cells were subsequently maintained in pH-adjusted PBS buffers ranging from pH 5.2 to 5.5 or 5.6 for 15 min. After aspiration of pH-adjusted PBS, infected cells were incubated in DMEM supplemented with 5% FBS for 3 h at 37°C. The cells were then fixed and stained using Hema 3 Fixative and Solutions (Fisher Scientific). Photomicrographs of cells containing or lacking syncytia were recorded using a light microscope (Russier et al., 2016, Reed et al., 2010). A baseline of no virus-induced syncytia formation was obtained by exposing the cells to pH 5.5 media, under which Vero cells formed no visible syncytia. Micrographs were scored positive for syncytia formation if a field contained at least two syncytia that had at least five nuclei. HA activation pH values for syncytia assays were reported as the highest pH that induced syncytia as judged by positive scoring.

### Polymerase activity assay

A dual-luciferase reporter assay system (Promega, Madison, WI, USA) was used to compare the polymerase activities of viral RNP complexes(Xu et al., 2016). The indicated viruses’ PB2, PB1, PA, and NP gene segments were separately cloned into the pCDNA3.1 expression plasmid. Mutations were introduced into the PB1 gene by site-directed mutagenesis (Invitrogen) according to the manufacturer’s protocol. All constructs were sequenced to ensure the absence of unwanted mutations. The primer sequences used for cloning are available upon request. The PB2, PB1, PA, and NP plasmids (125 ng each plasmid) were used to transfect 293T cells with the Luci luciferase reporter plasmid (10 ng) and the renilla internal control plasmid (2.5 ng). Cultures were incubated at 33°C and 37°C. Cell lysates were analyzed 24 h after transfection to measure firefly and renilla luciferase activities using GloMax 96 microplate luminometer (Promega).

### Statistical analyses

Differences between experimental groups were assessed using analysis of variance (ANOVA). *P* < 0.05 was considered to indicate a statistically significant difference.

### Nucleotide sequence accession numbers

The nucleotide sequences of 66 H3N2 CIVs isolated in this study are available in GenBank under accession numbers:ON877531-ON878058.

## Acknowledgments

This work was supported by the National Natural Science Foundation of China (32172838 and 32192451) and the 111 Project.

## Supplementary Tables and Figures

**Table S1.**
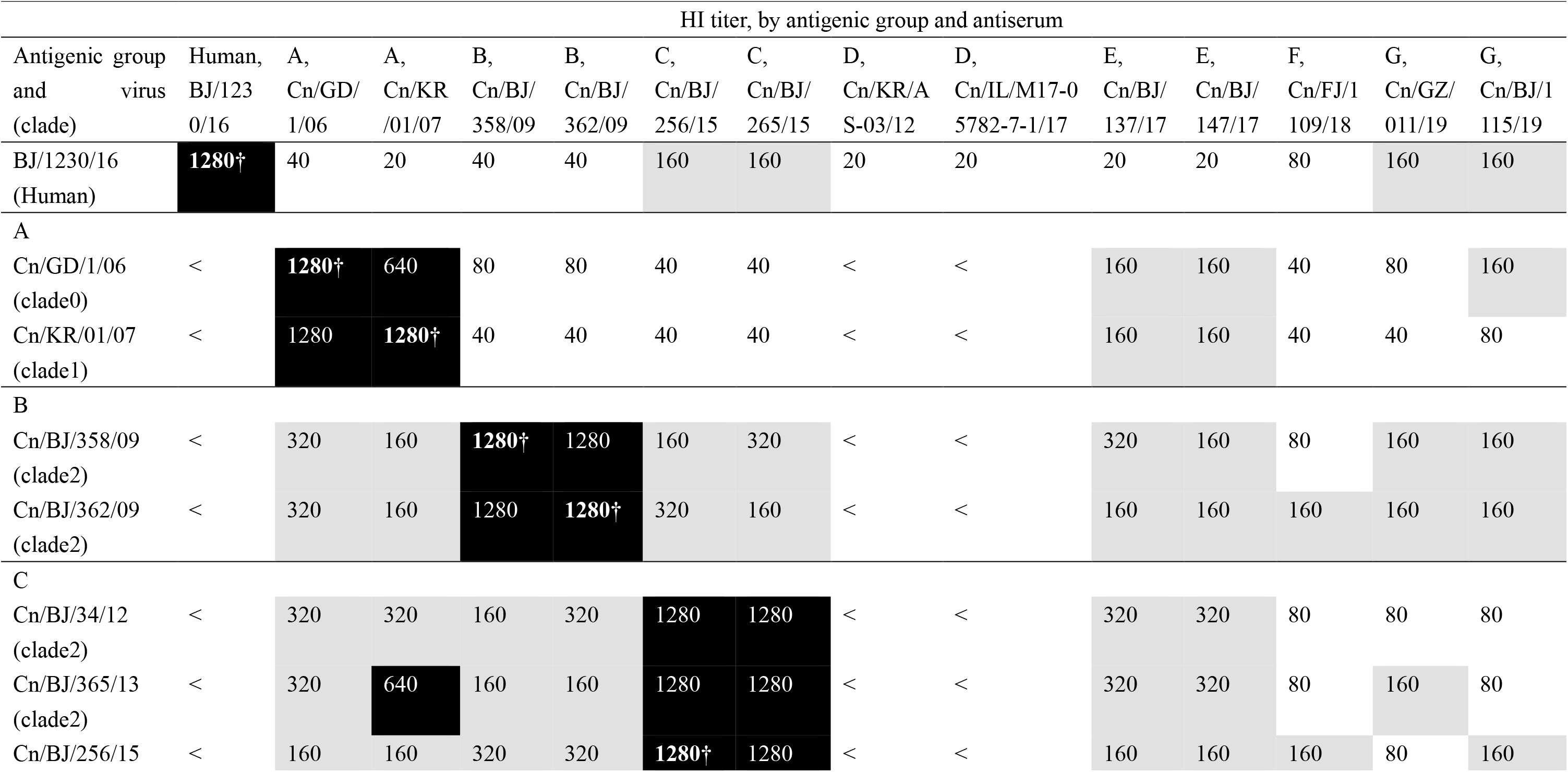
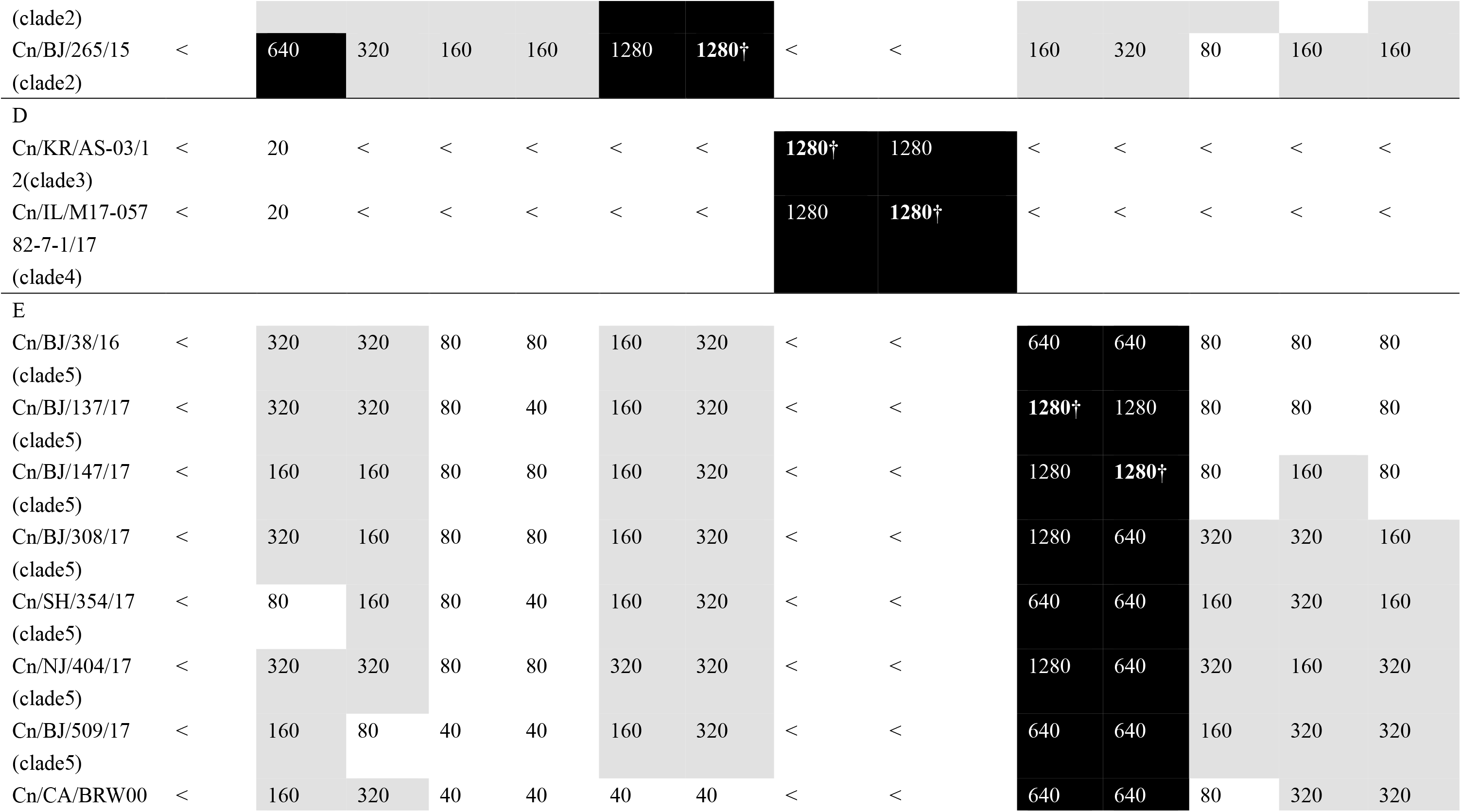
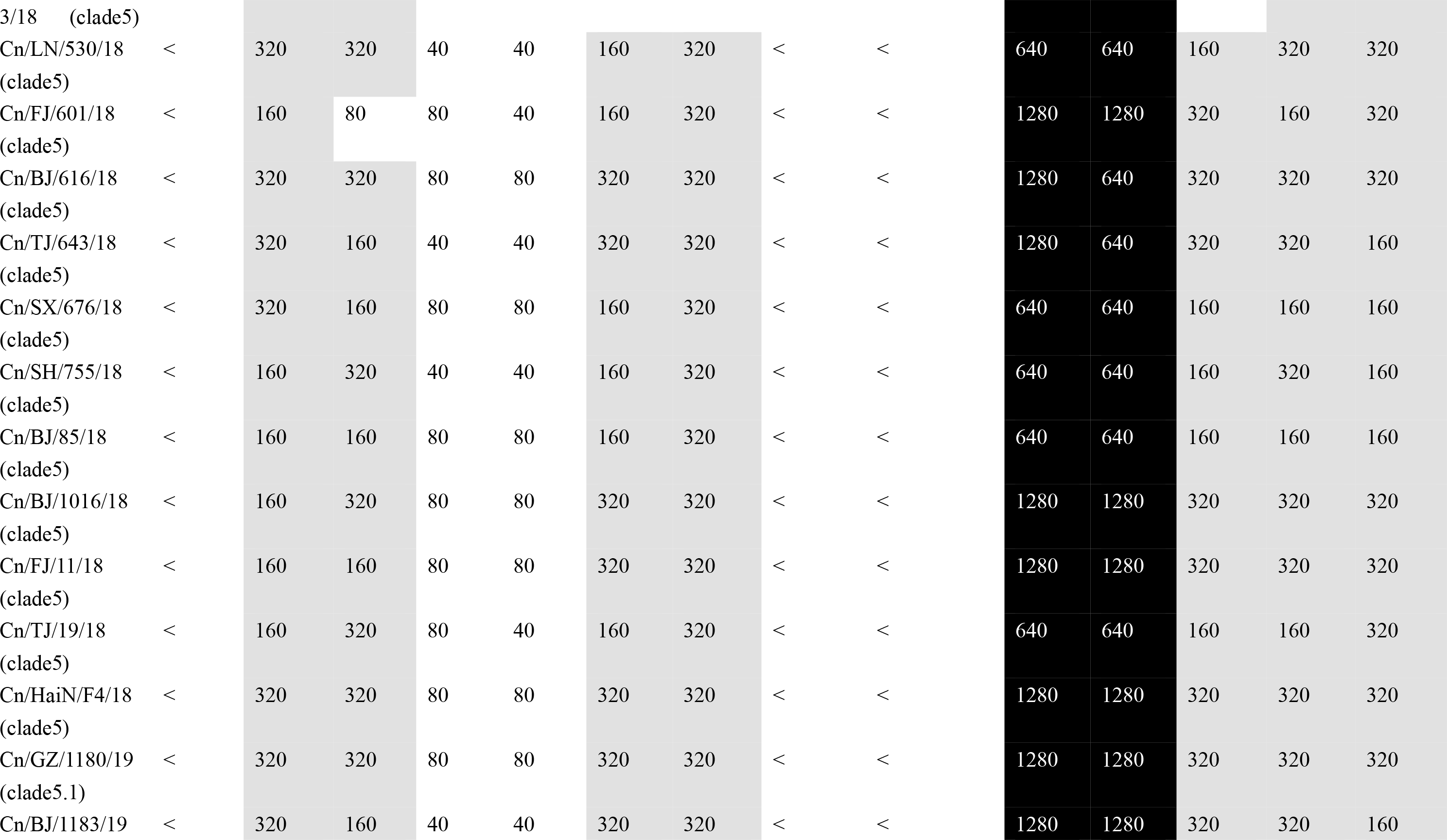
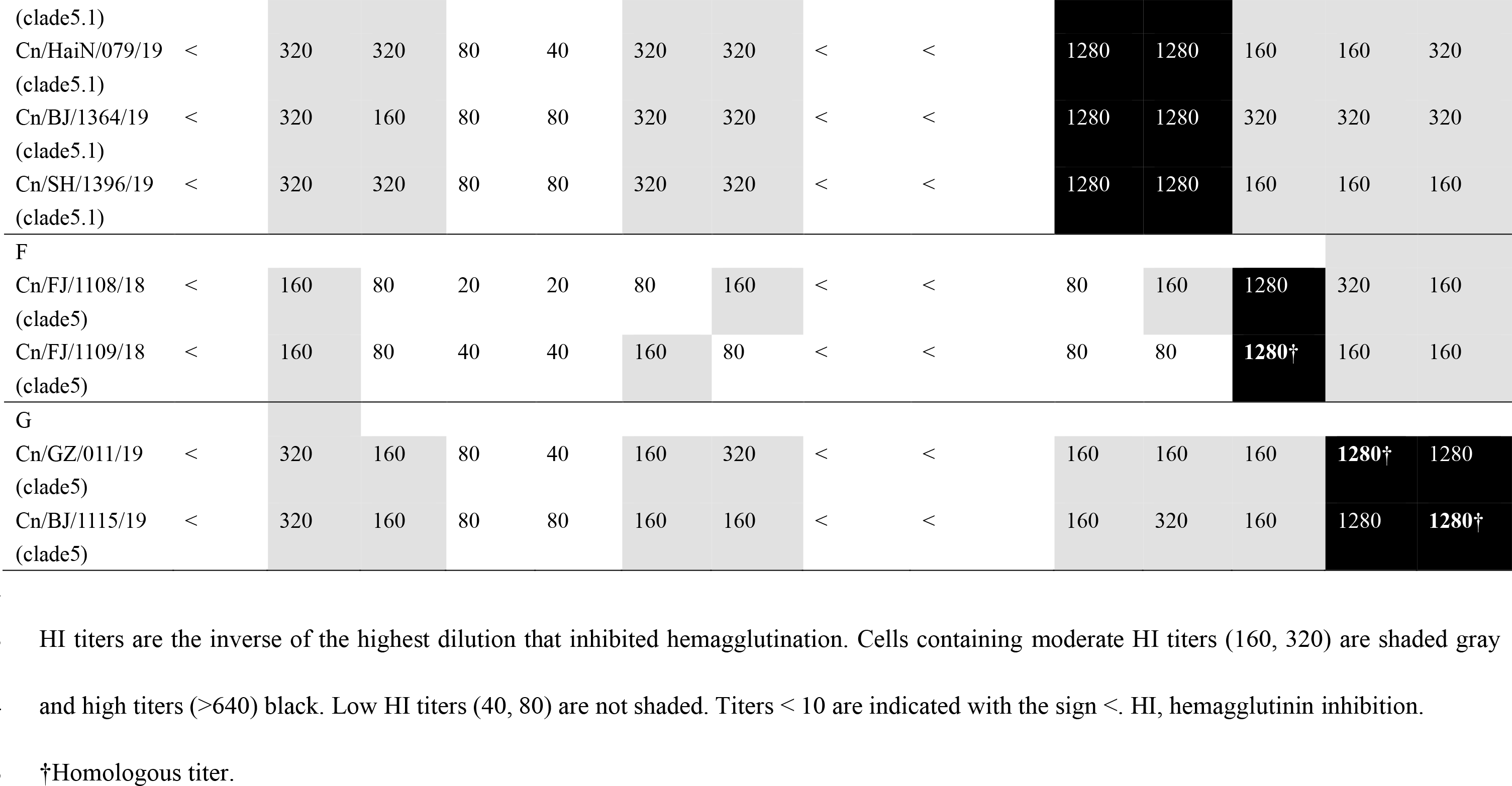
Antigenic analysis of H3N2 subtype canine influenza viruses in world.

| HI titer, by antigenic group and antiserum |  |  |  |  |  |  |  |  |  |  |  |  |  |  |
| --- | --- | --- | --- | --- | --- | --- | --- | --- | --- | --- | --- | --- | --- | --- |
| Antigenic group and virus (clade) | Human, BJ/123 0/16 | A, Cn/GD/1/06 | A, Cn/KR/01/07 | B, Cn/BJ/358/09 | B, Cn/BJ/362/09 | C, Cn/BJ/256/15 | C, Cn/BJ/265/15 | D, Cn/KR/A S-03/12 | D, Cn/IL/M17-0 5782-7-1/17 | E, Cn/BJ/137/17 | E, Cn/BJ/147/17 | F, Cn/FJ/1 109/18 | G, Cn/GZ/011/19 | G, Cn/BJ/1 115/19 |
| BJ/1230/16 (Human) | 1280† | 40 | 20 | 40 | 40 | 160 | 160 | 20 | 20 | 20 | 20 | 80 | 160 | 160 |
| A |  |  |  |  |  |  |  |  |  |  |  |  |  |  |
| Cn/GD/1/06 (clade0) | < | 1280† | 640 | 80 | 80 | 40 | 40 | < | < | 160 | 160 | 40 | 80 | 160 |
| Cn/KR/01/07 (clade1) | < | 1280 | 1280† | 40 | 40 | 40 | 40 | < | < | 160 | 160 | 40 | 40 | 80 |
| B |  |  |  |  |  |  |  |  |  |  |  |  |  |  |
| Cn/BJ/358/09 (clade2) | < | 320 | 160 | 1280† | 1280 | 160 | 320 | < | < | 320 | 160 | 80 | 160 | 160 |
| Cn/BJ/362/09 (clade2) | < | 320 | 160 | 1280 | 1280† | 320 | 160 | < | < | 160 | 160 | 160 | 160 | 160 |
| C |  |  |  |  |  |  |  |  |  |  |  |  |  |  |
| Cn/BJ/34/12 (clade2) | < | 320 | 320 | 160 | 320 | 1280 | 1280 | < | < | 320 | 320 | 80 | 80 | 80 |
| Cn/BJ/365/13 (clade2) | < | 320 | 640 | 160 | 160 | 1280 | 1280 | < | < | 320 | 320 | 80 | 160 | 80 |
| Cn/BJ/256/15 | < | 160 | 160 | 320 | 320 | 1280† | 1280 | < | < | 160 | 160 | 160 | 80 | 160 |
| (clade2) |  |  |  |  |  |  |  |  |  |  |  |  |  |  |
| Cn/BJ/265/15 | < | 640 | 320 | 160 | 160 | 1280 | 1280† | < | < | 160 | 320 | 80 | 160 | 160 |
| (clade2) |  |  |  |  |  |  |  |  |  |  |  |  |  |  |
| D |  |  |  |  |  |  |  |  |  |  |  |  |  |  |
| Cn/KR/AS-03/1 | < | 20 | < | < | < | < | < | 1280† | 1280 | < | < | < | < | < |
| 2(clade3) |  |  |  |  |  |  |  |  |  |  |  |  |  |  |
| Cn/IL/M17-057 | < | 20 | < | < | < | < | < | 1280 | 1280† | < | < | < | < | < |
| 82-7-1/17 |  |  |  |  |  |  |  |  |  |  |  |  |  |  |
| (clade4) |  |  |  |  |  |  |  |  |  |  |  |  |  |  |
| E |  |  |  |  |  |  |  |  |  |  |  |  |  |  |
| Cn/BJ/38/16 | < | 320 | 320 | 80 | 80 | 160 | 320 | < | < | 640 | 640 | 80 | 80 | 80 |
| (clade5) |  |  |  |  |  |  |  |  |  |  |  |  |  |  |
| Cn/BJ/137/17 | < | 320 | 320 | 80 | 40 | 160 | 320 | < | < | 1280† | 1280 | 80 | 80 | 80 |
| (clade5) |  |  |  |  |  |  |  |  |  |  |  |  |  |  |
| Cn/BJ/147/17 | < | 160 | 160 | 80 | 80 | 160 | 320 | < | < | 1280 | 1280† | 80 | 160 | 80 |
| (clade5) |  |  |  |  |  |  |  |  |  |  |  |  |  |  |
| Cn/BJ/308/17 | < | 320 | 160 | 80 | 80 | 160 | 320 | < | < | 1280 | 640 | 320 | 320 | 160 |
| (clade5) |  |  |  |  |  |  |  |  |  |  |  |  |  |  |
| Cn/SH/354/17 | < | 80 | 160 | 80 | 40 | 160 | 320 | < | < | 640 | 640 | 160 | 320 | 160 |
| (clade5) |  |  |  |  |  |  |  |  |  |  |  |  |  |  |
| Cn/NJ/404/17 | < | 320 | 320 | 80 | 80 | 320 | 320 | < | < | 1280 | 640 | 320 | 160 | 320 |
| (clade5) |  |  |  |  |  |  |  |  |  |  |  |  |  |  |
| Cn/BJ/509/17 | < | 160 | 80 | 40 | 40 | 160 | 320 | < | < | 640 | 640 | 160 | 320 | 320 |
| (clade5) |  |  |  |  |  |  |  |  |  |  |  |  |  |  |
| Cn/CA/BRW00 | < | 160 | 320 | 40 | 40 | 40 | 40 | < | < | 640 | 640 | 80 | 320 | 320 |
| 3/18 (clade5) |  |  |  |  |  |  |  |  |  |  |  |  |  |  |
| Cn/LN/530/18 (clade5) | < | 320 | 320 | 40 | 40 | 160 | 320 | < | < | 640 | 640 | 160 | 320 | 320 |
| Cn/FJ/601/18 (clade5) | < | 160 | 80 | 80 | 40 | 160 | 320 | < | < | 1280 | 1280 | 320 | 160 | 320 |
| Cn/BJ/616/18 (clade5) | < | 320 | 320 | 80 | 80 | 320 | 320 | < | < | 1280 | 640 | 320 | 320 | 320 |
| Cn/TJ/643/18 (clade5) | < | 320 | 160 | 40 | 40 | 320 | 320 | < | < | 1280 | 640 | 320 | 320 | 160 |
| Cn/SX/676/18 (clade5) | < | 320 | 160 | 80 | 80 | 160 | 320 | < | < | 640 | 640 | 160 | 160 | 160 |
| Cn/SH/755/18 (clade5) | < | 160 | 320 | 40 | 40 | 160 | 320 | < | < | 640 | 640 | 160 | 320 | 160 |
| Cn/BJ/85/18 (clade5) | < | 160 | 160 | 80 | 80 | 160 | 320 | < | < | 640 | 640 | 160 | 160 | 160 |
| Cn/BJ/1016/18 (clade5) | < | 160 | 320 | 80 | 80 | 320 | 320 | < | < | 1280 | 1280 | 320 | 320 | 320 |
| Cn/FJ/11/18 (clade5) | < | 160 | 160 | 80 | 80 | 320 | 320 | < | < | 1280 | 1280 | 320 | 320 | 320 |
| Cn/TJ/19/18 (clade5) | < | 160 | 320 | 80 | 40 | 160 | 320 | < | < | 640 | 640 | 160 | 160 | 320 |
| Cn/HaiN/F4/18 (clade5) | < | 320 | 320 | 80 | 80 | 320 | 320 | < | < | 1280 | 1280 | 320 | 320 | 320 |
| Cn/GZ/1180/19 (clade5.1) | < | 320 | 320 | 80 | 80 | 320 | 320 | < | < | 1280 | 1280 | 320 | 320 | 320 |
| Cn/BJ/1183/19 | < | 320 | 160 | 40 | 40 | 320 | 320 | < | < | 1280 | 1280 | 320 | 320 | 160 |
| (clade5.1) |  |  |  |  |  |  |  |  |  |  |  |  |  |  |
| Cn/HaiN/079/19 | < | 320 | 320 | 80 | 40 | 320 | 320 | < | < | 1280 | 1280 | 160 | 160 | 320 |
| (clade5.1) |  |  |  |  |  |  |  |  |  |  |  |  |  |  |
| Cn/BJ/1364/19 | < | 320 | 160 | 80 | 80 | 320 | 320 | < | < | 1280 | 1280 | 320 | 320 | 320 |
| (clade5.1) |  |  |  |  |  |  |  |  |  |  |  |  |  |  |
| Cn/SH/1396/19 | < | 320 | 320 | 80 | 80 | 320 | 320 | < | < | 1280 | 1280 | 160 | 160 | 160 |
| (clade5.1) |  |  |  |  |  |  |  |  |  |  |  |  |  |  |
| F |  |  |  |  |  |  |  |  |  |  |  |  |  |  |
| Cn/FJ/1108/18 | < | 160 | 80 | 20 | 20 | 80 | 160 | < | < | 80 | 160 | 1280 | 320 | 160 |
| (clade5) |  |  |  |  |  |  |  |  |  |  |  |  |  |  |
| Cn/FJ/1109/18 | < | 160 | 80 | 40 | 40 | 160 | 80 | < | < | 80 | 80 | 1280† | 160 | 160 |
| (clade5) |  |  |  |  |  |  |  |  |  |  |  |  |  |  |
| G |  |  |  |  |  |  |  |  |  |  |  |  |  |  |
| Cn/GZ/011/19 | < | 320 | 160 | 80 | 40 | 160 | 320 | < | < | 160 | 160 | 160 | 1280† | 1280 |
| (clade5) |  |  |  |  |  |  |  |  |  |  |  |  |  |  |
| Cn/BJ/1115/19 | < | 320 | 160 | 80 | 80 | 160 | 160 | < | < | 160 | 320 | 160 | 1280 | 1280† |
| (clade5) |  |  |  |  |  |  |  |  |  |  |  |  |  |  |

HI titers are the inverse of the highest dilution that inhibited hemagglutination. Cells containing moderate HI titers (160, 320) are shaded gray
and high titers (>640) black. Low HI titers (40, 80) are not shaded. Titers < 10 are indicated with the sign <. HI, hemagglutinin inhibition.
†Homologous titer.

**Table S2.**
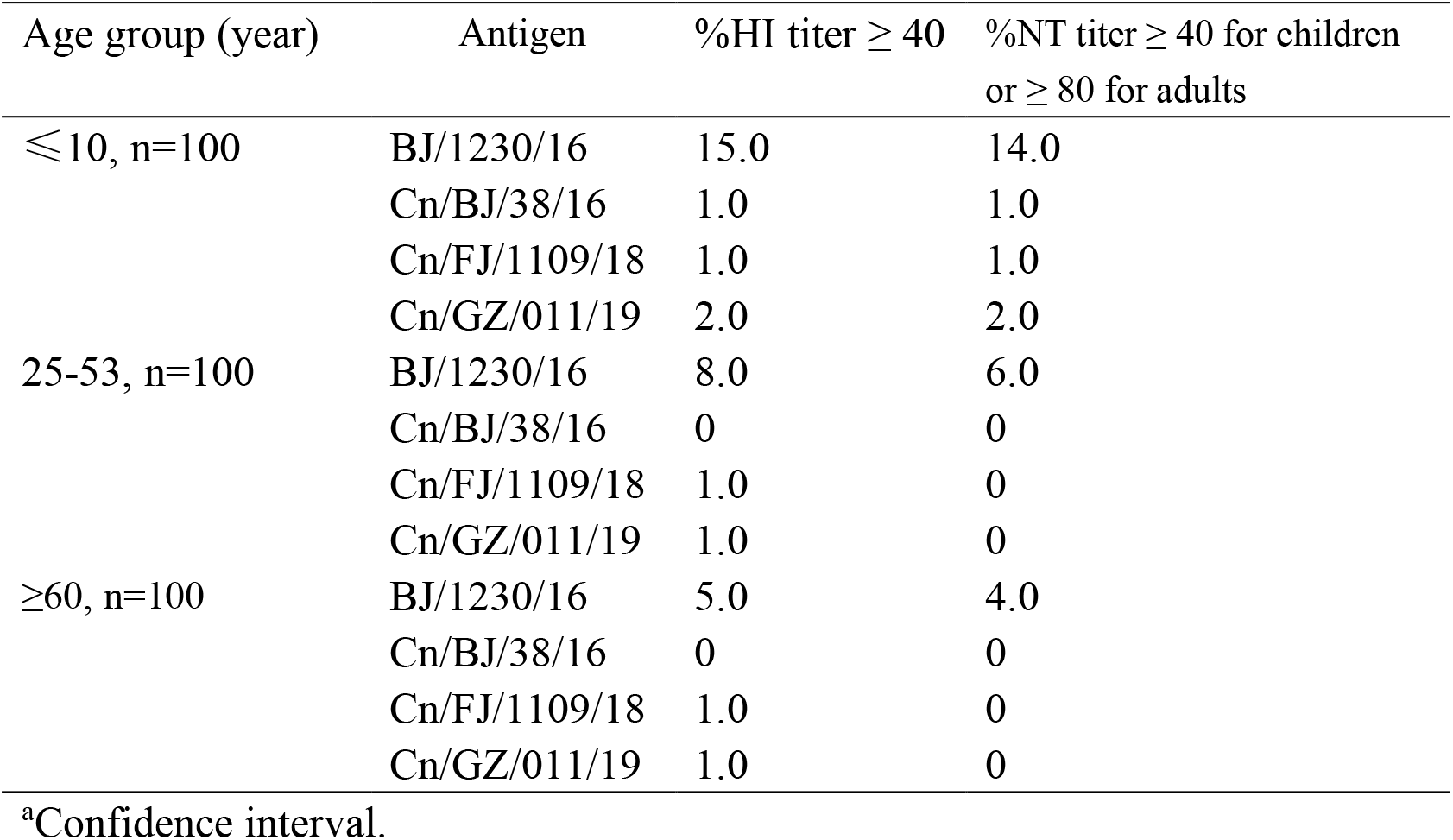
Cross-reactive antibody responses against influenza A (H3N2) viruses of human sera collected in China.

| Age group (year) | Antigen | %HI titer $\geq 40$ | %NT titer $\geq 40$ for children<br>or $\geq 80$ for adults |
| --- | --- | --- | --- |
| $\leq 10$, n=100 | BJ/1230/16 | 15.0 | 14.0 |
|  | Cn/BJ/38/16 | 1.0 | 1.0 |
|  | Cn/FJ/1109/18 | 1.0 | 1.0 |
|  | Cn/GZ/011/19 | 2.0 | 2.0 |
| 25-53, n=100 | BJ/1230/16 | 8.0 | 6.0 |
|  | Cn/BJ/38/16 | 0 | 0 |
|  | Cn/FJ/1109/18 | 1.0 | 0 |
|  | Cn/GZ/011/19 | 1.0 | 0 |
| $\geq 60$, n=100 | BJ/1230/16 | 5.0 | 4.0 |
|  | Cn/BJ/38/16 | 0 | 0 |
|  | Cn/FJ/1109/18 | 1.0 | 0 |
|  | Cn/GZ/011/19 | 1.0 | 0 |

<sup>a</sup>Confidence interval.

**Table S3.** Transmission of H3N2 reassortant viruses in dogs.

| Virus | Inoculated Animals |  | Contacted Animals |  |  |
| --- | --- | --- | --- | --- | --- |
|  | No. with virus detection (peak titer mean log<sub>10</sub>EID<sub>50</sub>/ml)‡ | No. with seroconversion (HI titer range)§ | Time of first infection (dpi) | No. with virus detection (peak titer mean log<sub>10</sub>EID<sub>50</sub>/ml)‡ | No. with seroconversion (HI titer range)§ |
| BJ15 | 3/3(4.17) | 3/3(320-640) | 4 | 3/3(3.58) | 3/3(160-320) |
| HaiN/19 | 3/3(6.17) | 3/3(1280) | 2 | 3/3(6.15) | 3/3(320-640) |
| rgHaiN19PB2 | 3/3(3.5) | 3/3(320) | 6 | 3/3(3.25) | 3/3(160-320) |
| rgHaiN19PB1 | 3/3(5.75) | 3/3(320-640) | 2 | 3/3(5.5) | 3/3(320) |
| rgHaiN19PA | 3/3(4.25) | 3/3(320) | 4 | 3/3(4.25) | 3/3(160-320) |
| rgHaiN19HA | 3/3(6.15) | 3/3(640) | 2 | 3/3(5.75) | 3/3(640) |
| rgHaiN19NP | 3/3(4.5) | 3/3(160-320) | 4 | 3/3(3.5) | 3/3(160) |
| rgHaiN19NA | 3/3(4.25) | 3/3(320) | 4 | 3/3(3.25) | 3/3(160-320) |
| rgHaiN19M | 3/3(4.75) | 3/3(320-640) | 4 | 3/3(3.5) | 3/3(160-320) |
| rgHaiN19NS | 3/3(4.35) | 3/3(320) | 4 | 3/3(4.2) | 3/3(160-320) |

‡Number of animals in which virus was detected in nasal swabs. Values in parentheses indicate nasal swab titers, which are expressed as mean log<sub>10</sub> peak virus titer observed within the first 9 dpi.
§Number of animals in which seroconversion was observed. Values in parentheses show HI antibody titers in sera collected at 21 dpi. HI titers were determined using the homologous virus as test antigen.

**Table S4.** Transmission of H3N2 reassortant viruses in ferrets.

| Virus | Inoculated Animals |  | Aerosol Animals |  |
| --- | --- | --- | --- | --- |
|  | No. with virus<br>detection | No. with<br>seroconversion | No. with virus<br>detection | No. with<br>seroconversion |
|  | (peak titer mean<br>log<sub>10</sub>EID<sub>50</sub>/ml)‡ | (HI titer range)§ | (peak titer mean<br>log<sub>10</sub>EID<sub>50</sub>/ml)‡ | (HI titer range)§ |
| BJ15 | 3/3(5.75) | 3/3(320-640) | 0/3 | 0/3 |
| HaiN/19 | 3/3(7.84) | 3/3(640-1280) | 3/3(5.75) | 3/3(320-640) |
| rgHaiN19PB2 | 3/3(3.75) | 3/3(320-640) | 0/3 | 0/3 |
| rgHaiN19PB1 | 3/3(6.83) | 3/3(640) | 2/3(4.25) | 2/3(160) |
| rgHaiN19PA | 3/3(4.75) | 3/3(320-640) | 0/3 | 0/3 |
| rgHaiN19HA | 3/3(7.25) | 3/3(160-640) | 2/3(4.75) | 2/3(80-160) |
| rgHaiN19NP | 3/3(6.15) | 3/3(160-320) | 1/3(2.75) | 1/3(40) |
| rgHaiN19NA | 3/3(4.75) | 3/3(160-320) | 0/3 | 0/3 |
| rgHaiN19M | 3/3(4.5) | 3/3(160-320) | 0/3 | 0/3 |
| rgHaiN19NS | 3/3(4.25) | 3/3(160-320) | 0/3 | 0/3 |

‡Number of animals in which virus was detected in nasal washes. Values in parentheses indicate nasal wash titers, which are expressed as mean log<sub>10</sub> peak virus titer observed within the first 9 dpi.
§Number of animals in which seroconversion was observed. Values in parentheses show HI antibody titers in sera collected at 21 dpi. HI titers were determined using the homologous virus as test antigen.

**Fig S1.**
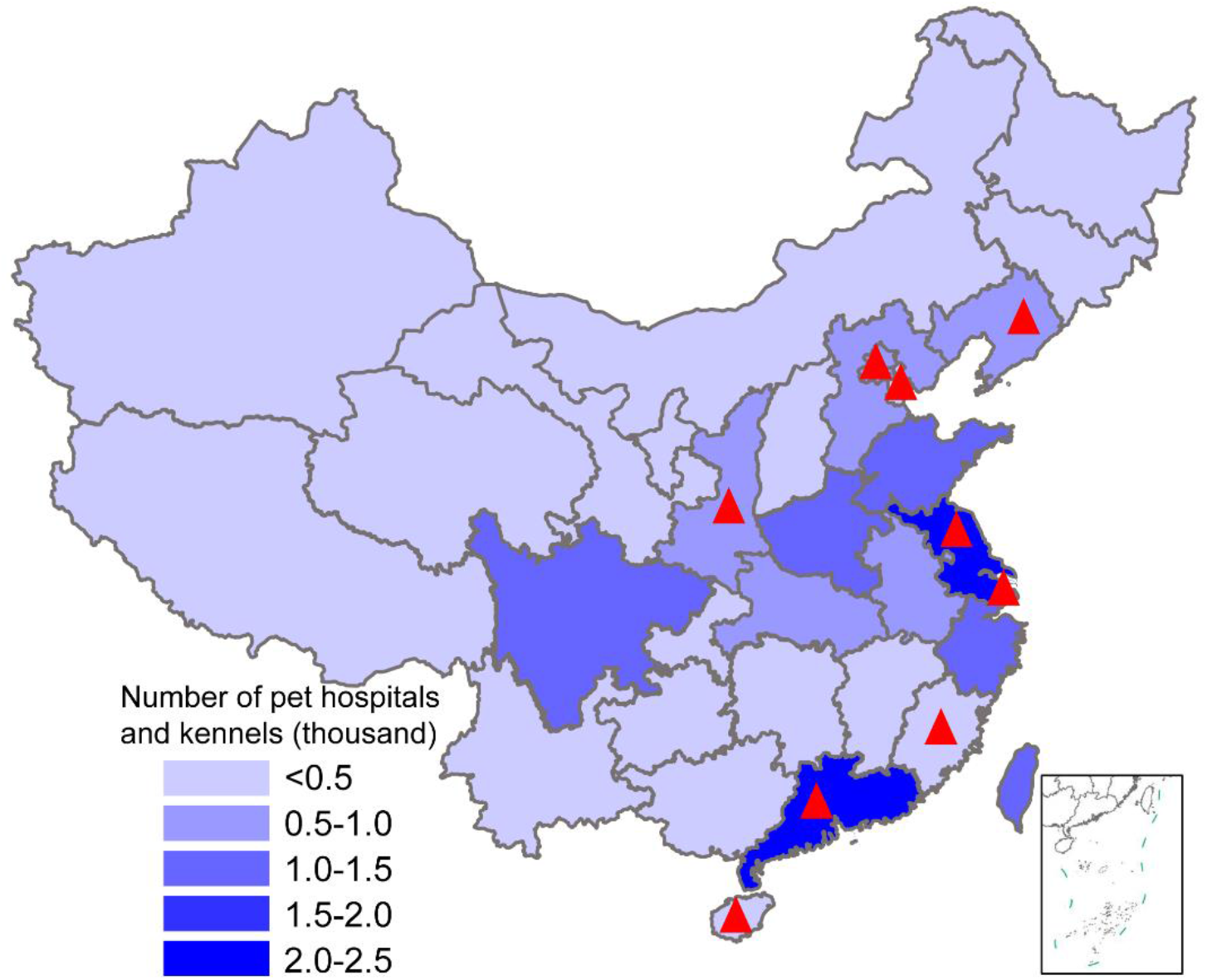
Map showing pet hospital and kennel density in China and geographic location of influenza surveillance in dogs from 2012 to 2019. The number of pet hospitals and kennels in each province is the average number from 2015 to 2019; data were from the National Bureau of Statistics of China. Red triangles indicate provinces where surveillance was conducted.

**Fig S2.**
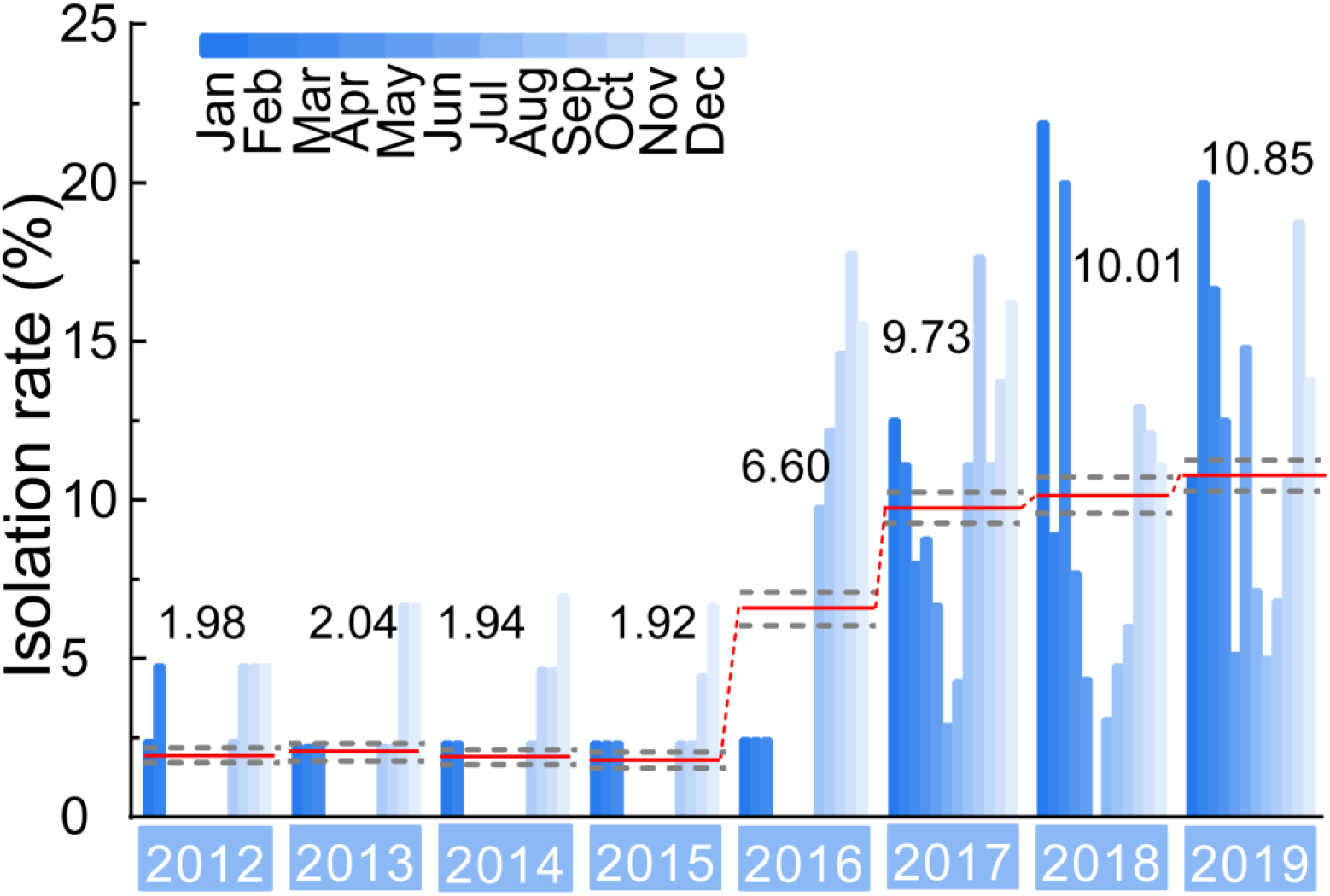
Isolation rate (%) of H3N2 CIVs during 2012–2019 from dogs with respiratory symptoms. Histogram indicates monthly isolation rates; red horizontal line (connected with dots) and the numbers above indicates mean annual isolation rates; and gray horizontal dashed line indicates 95% CI.

**Fig S3.**
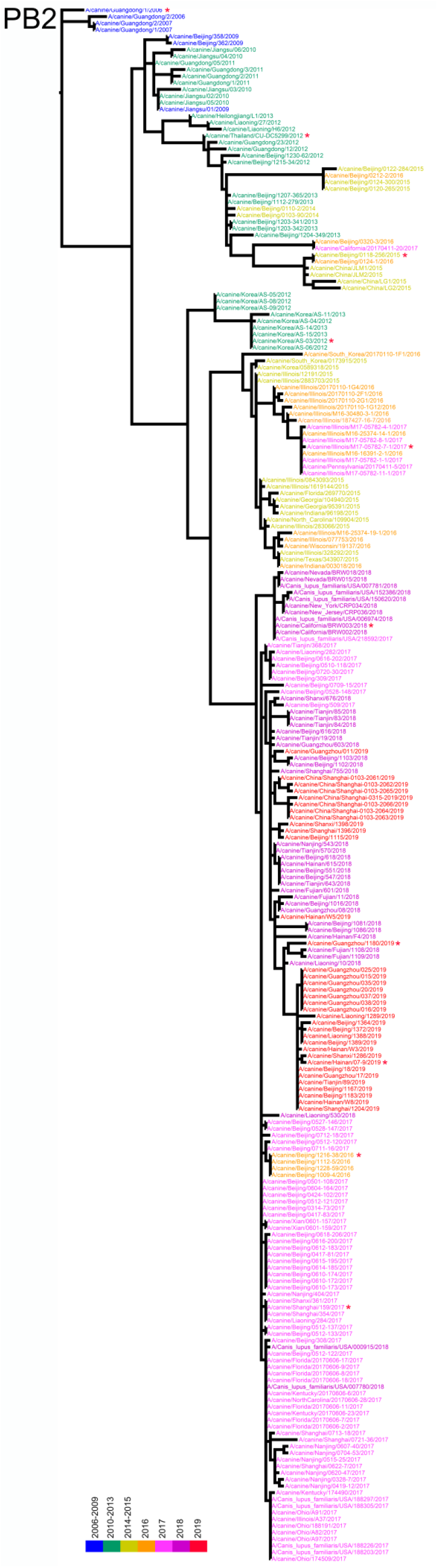
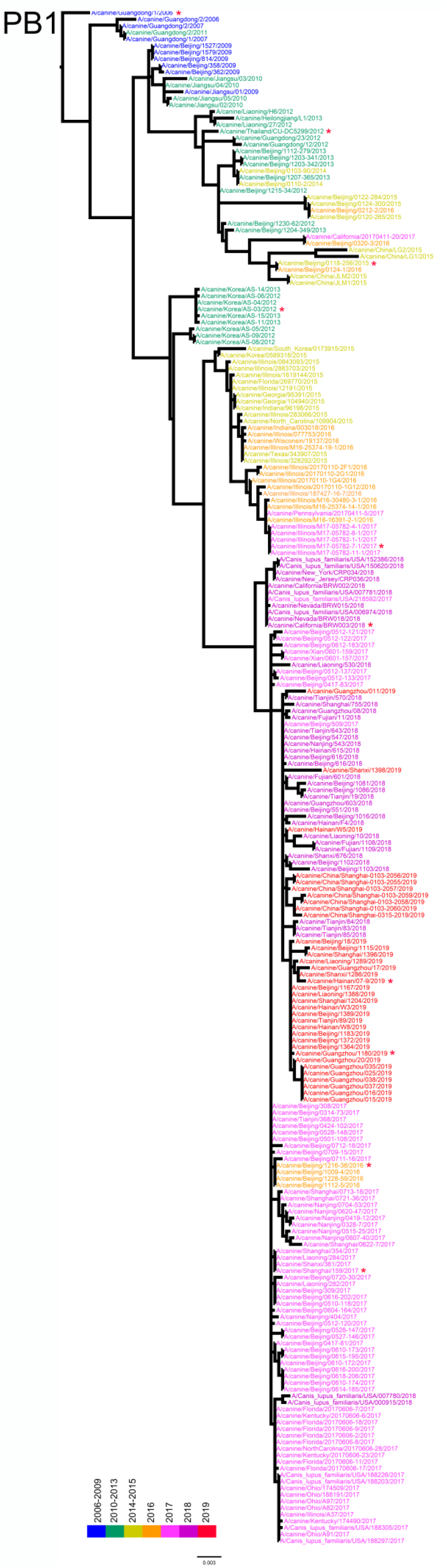
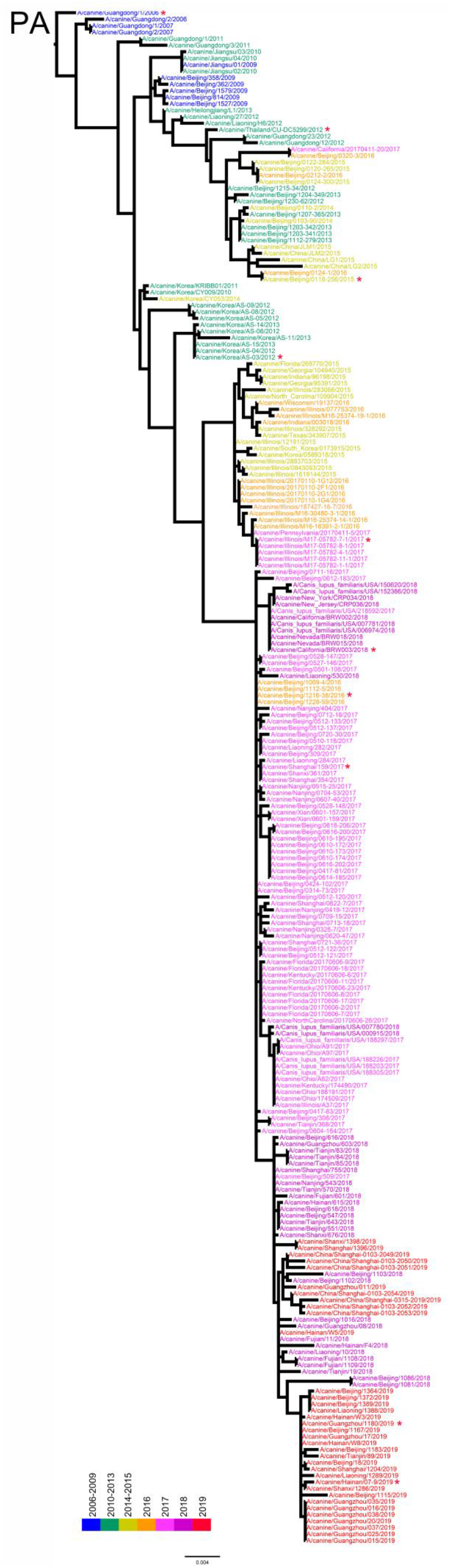
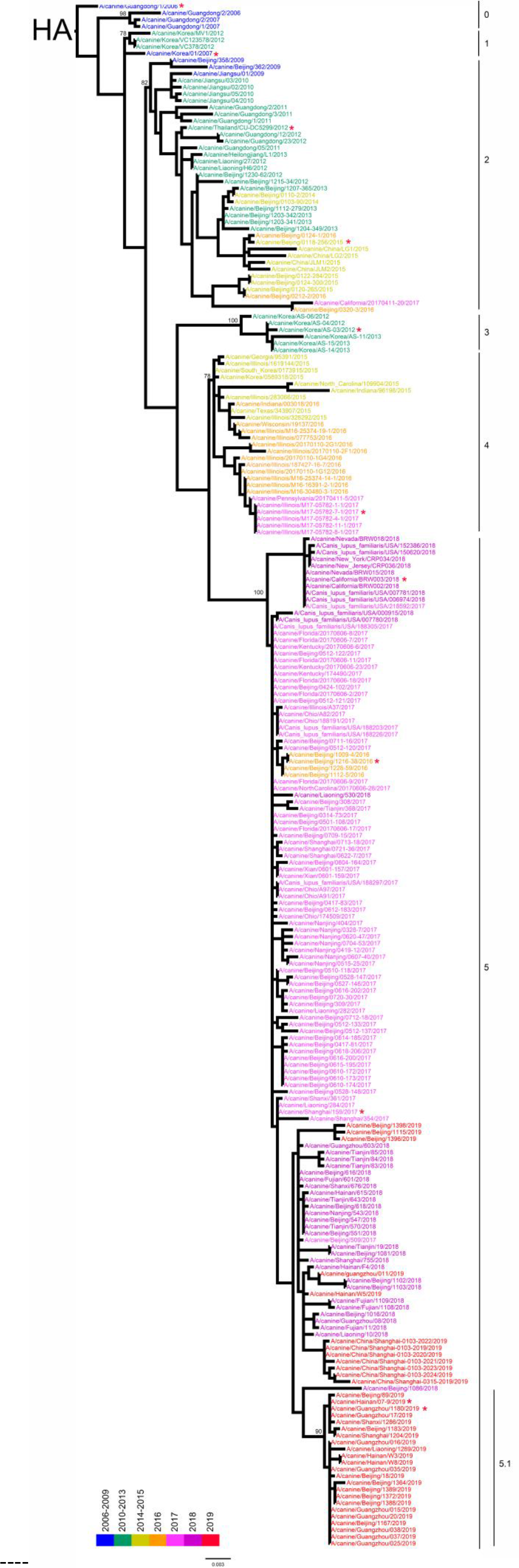
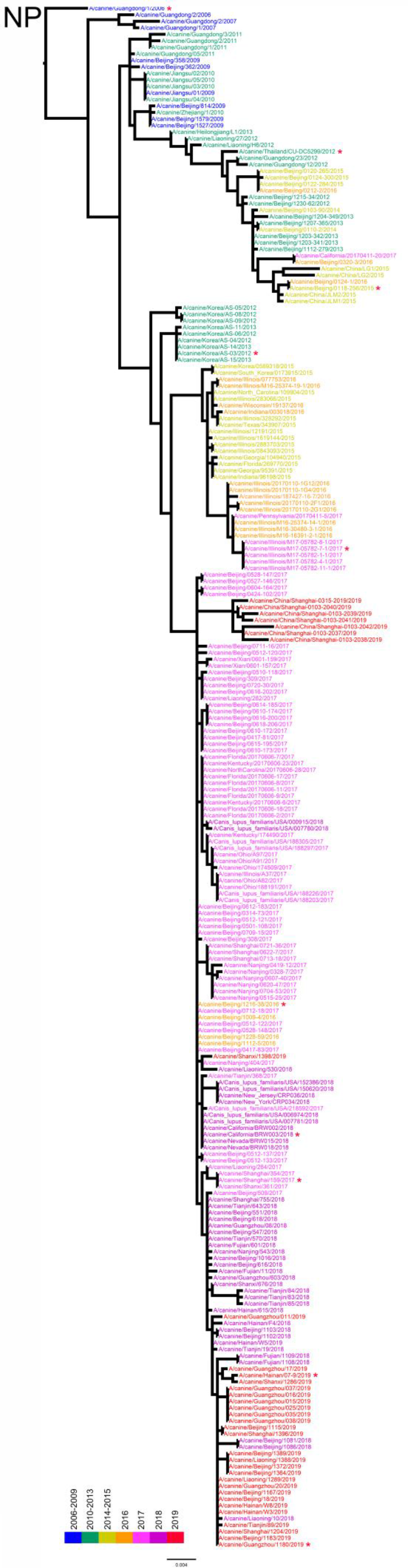
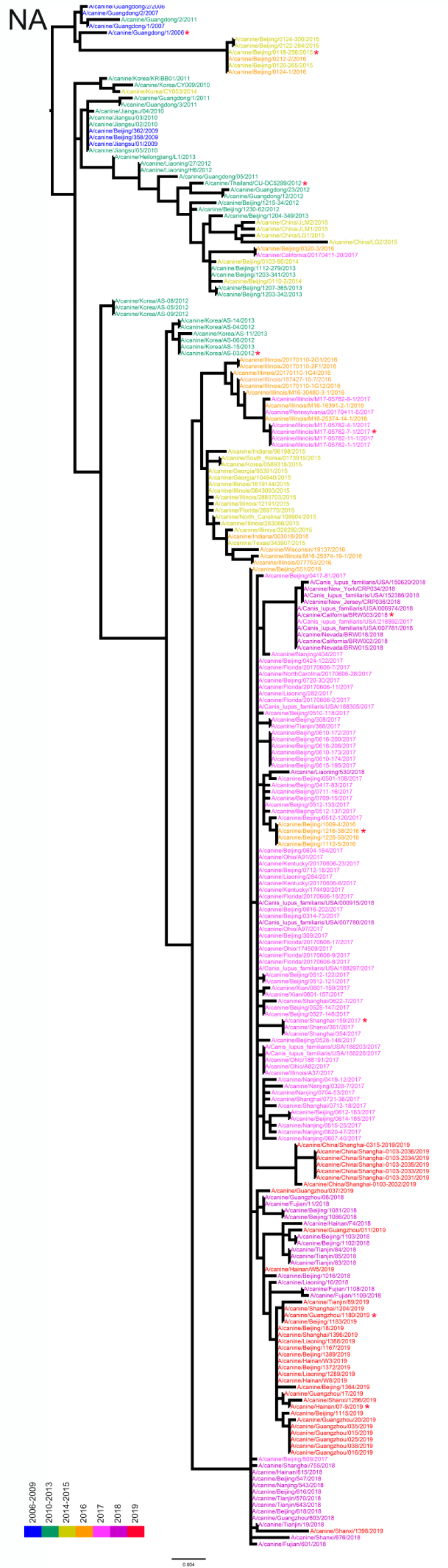
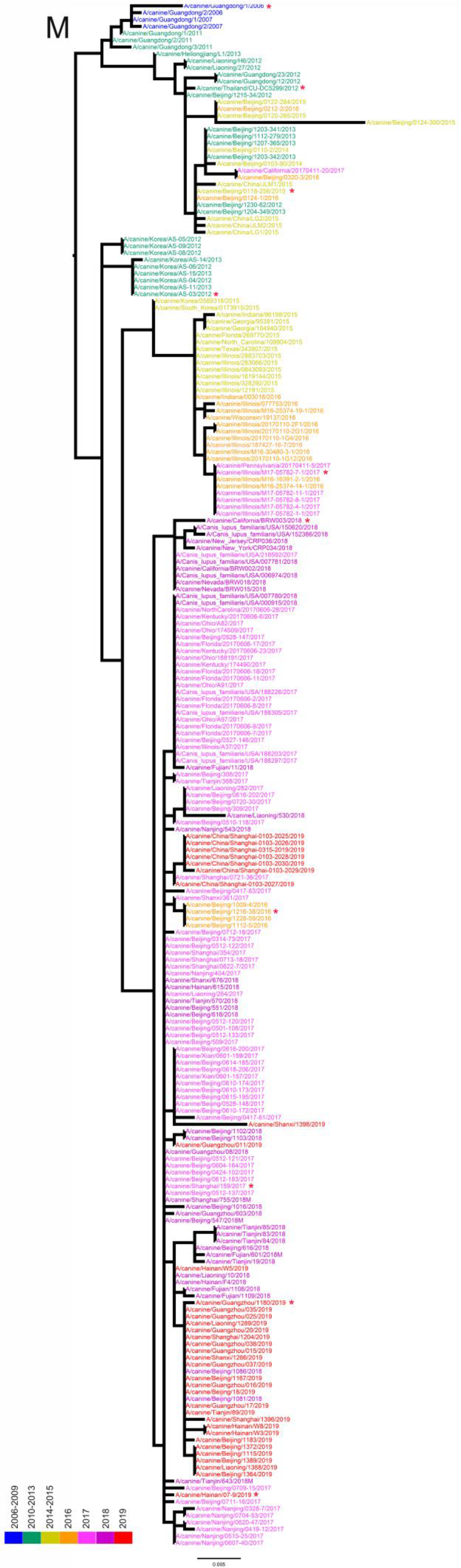
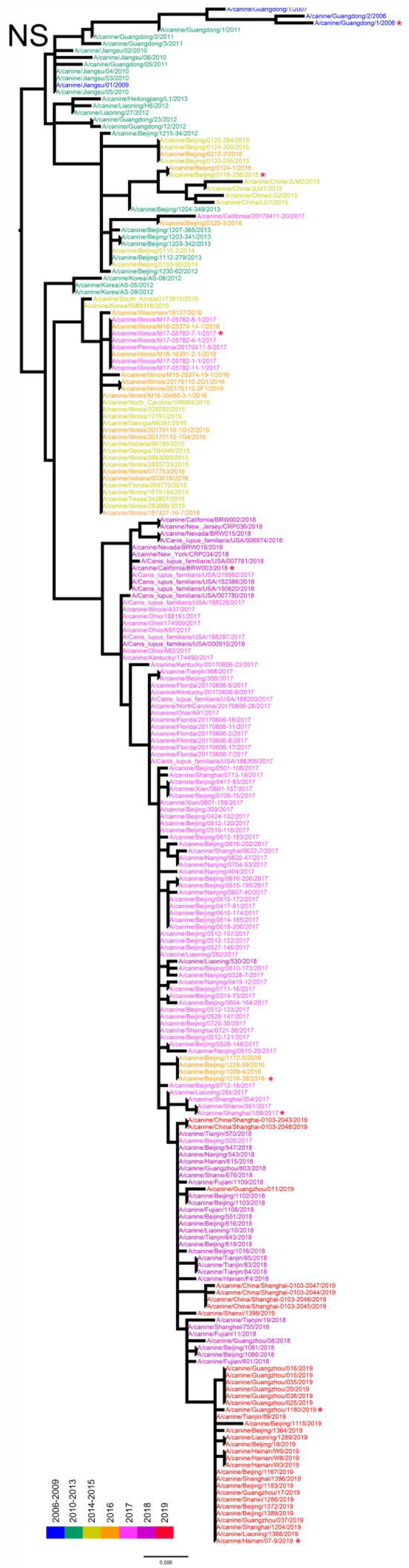
Phylogenetic relationships of fully sequenced H3N2 CIVs worldwide from 2006 to 2019. Phylogenetic trees were estimated using genetic distances calculated by maximum likelihood under the GTRGAMMA + I model. The color of strain names indicates year of isolation (see color bar).Scale bar is in units of nucleotide substitutions per site. Node labels in HA gene tree represent bootstrap values. Red asterisks indicate the phylogenetic position of H3N2 CIVs used in animal experiments.

**Fig S4.**
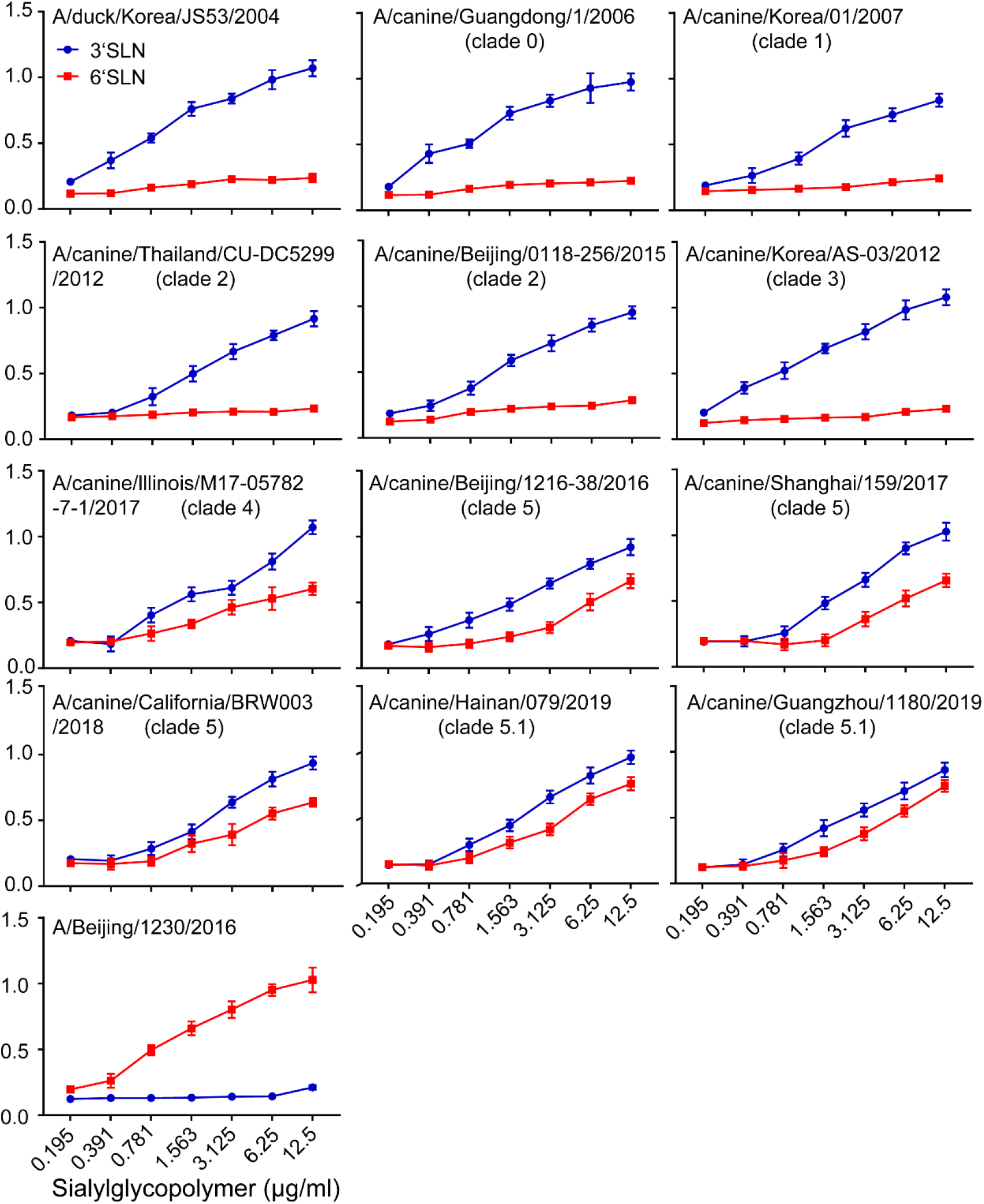
Characterization of the receptor-binding properties of H3N2 CIVs. The direct binding of the virus to sialylglycopolymers containing either 2,3-linked (blue) or 2,6-linked (red) sialic acids was tested.

**Fig S5.**
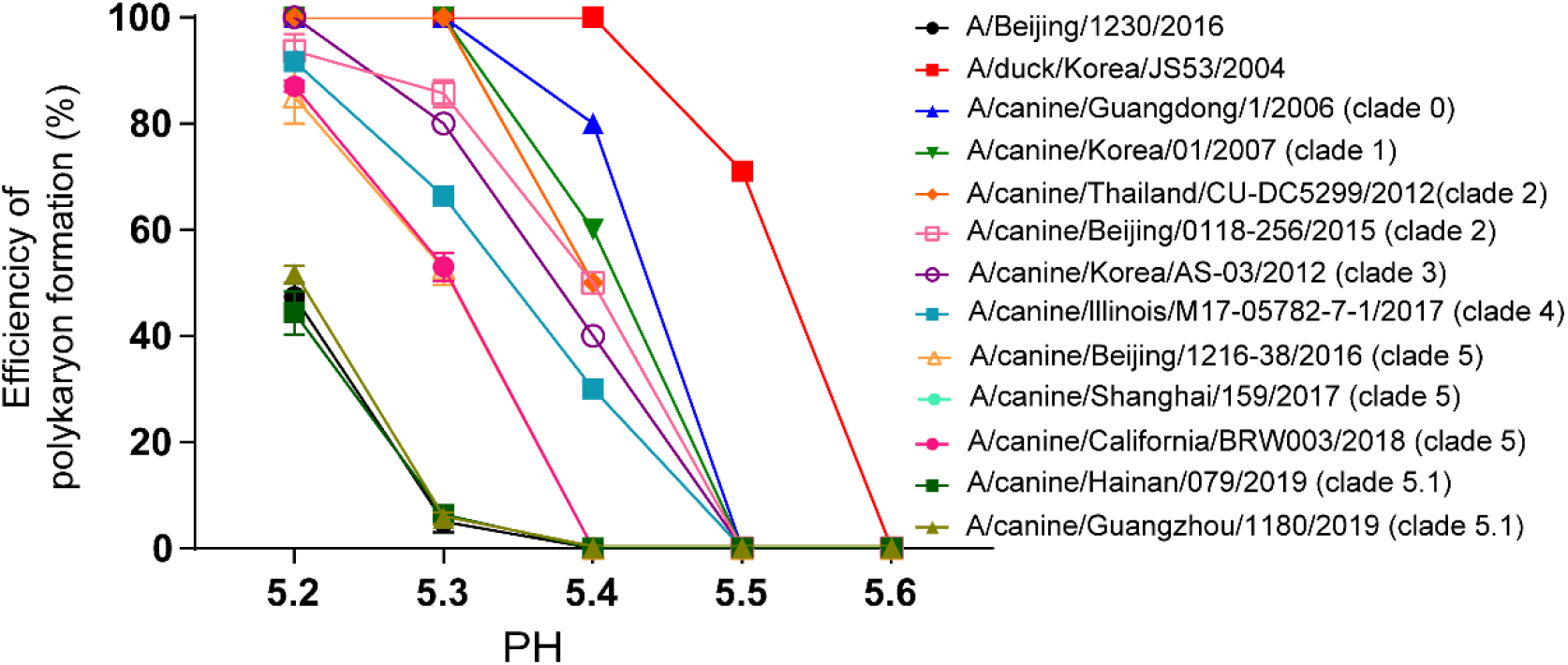
Syncytia assay results for H3N2 CIVs. The efficiency of polykaryon formation over a pH range of 5.2–5.6 was estimated from the number of nuclei in the polykaryons divided by the total number of nuclei in the same field. The mean and standard deviation determined from five randomly chosen fields of cell culture are shown.

**Fig S6.**
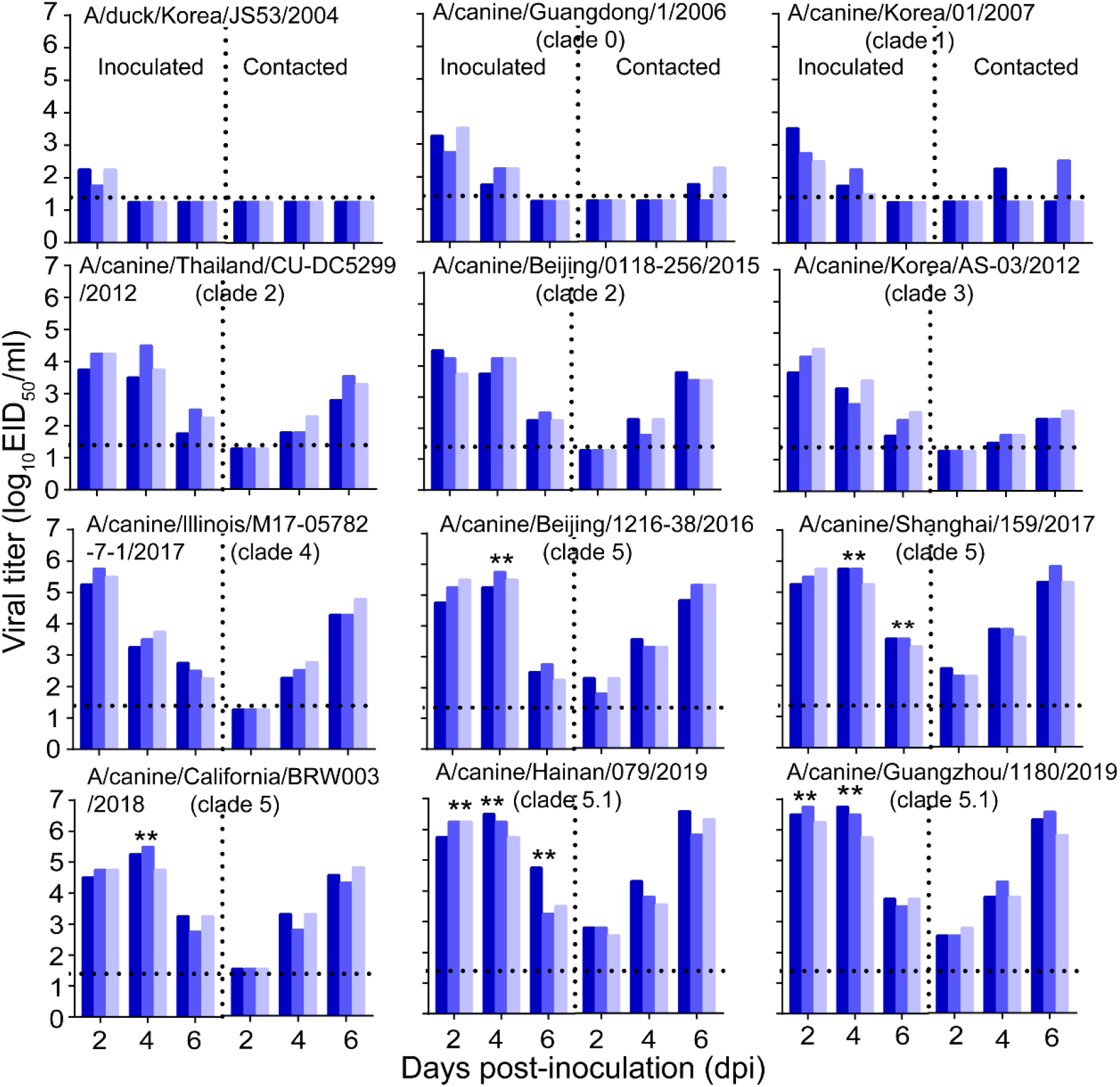
Direct contact transmission of H3N2 CIVs in dogs. Groups of three female beagles housed in a cage placed inside an isolator were inoculated intranasally with 10^6^ EID_50_ of indicated virus. Twenty-four hours later, the three inoculated animals were individually paired by co-housing with a direct-contact dog. Nasal swabs were collected every other day from all animals for virus shedding detection from day 2 of the initial infection. Each color bar represents the virus titer of an individual animal. No data are displayed when the virus was not detected from all of the groups. Dashed lines indicate the lower limit of virus detection. Statistical significance of clade 5.1 or clade 5 viruses relative to other viruses in the inoculated animals were assessed using two-way ANOVA (**, *P* < 0.01).

**Fig S7.**
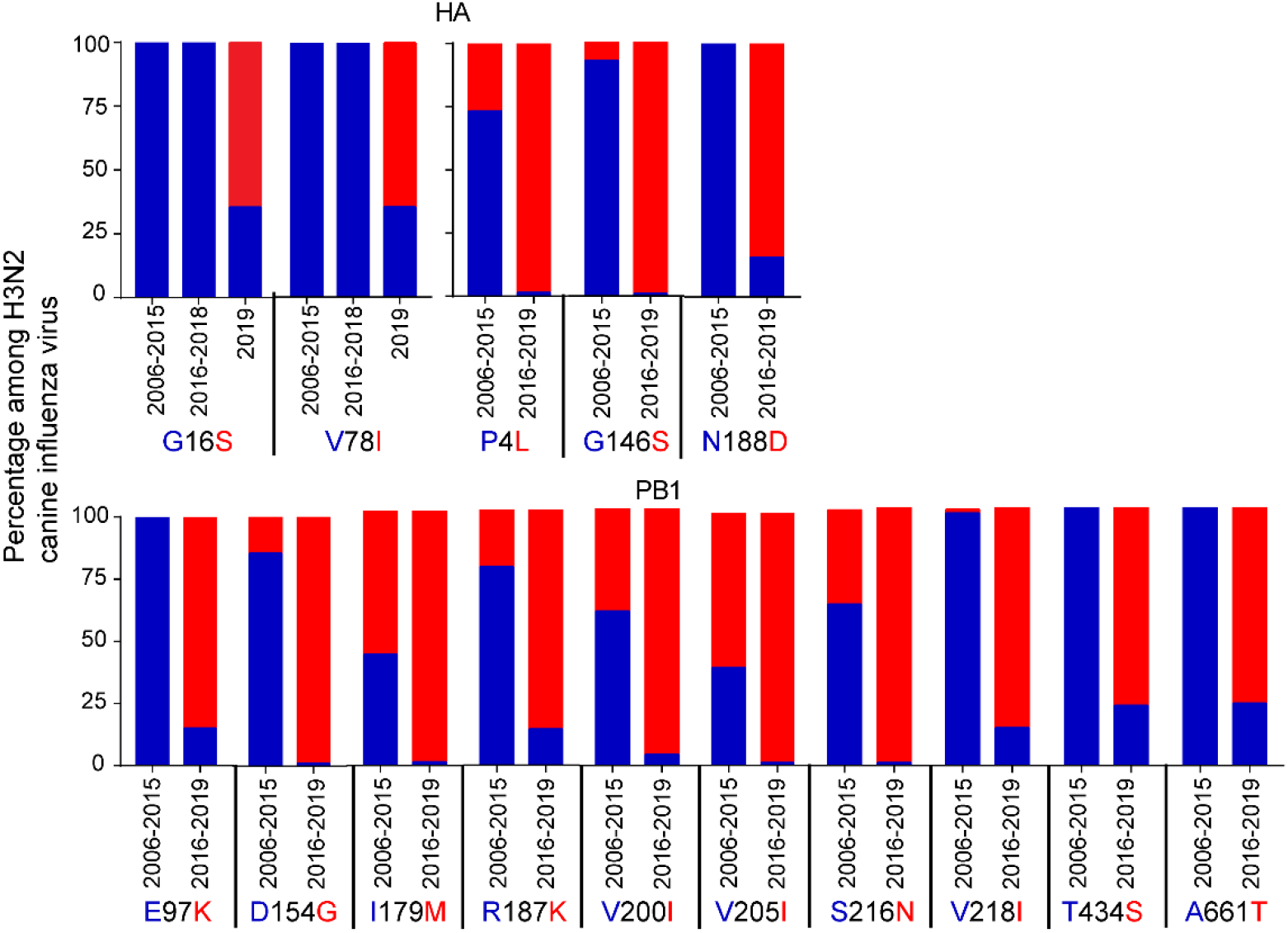
Detection frequency of the 15 amino acids that differed between 2006-2015 H3N2 CIVs and 2016-2019 H3N2 CIVs among HA (n=298) and PB1(n=298). Amino acid residues are colored in blue or red.

